# Borg tandem repeats undergo rapid evolution and are under strong selection to create new intrinsically disordered regions in proteins

**DOI:** 10.1101/2022.05.18.492195

**Authors:** Marie C. Schoelmerich, Rohan Sachdeva, Lucas Waldburger, Jacob West-Roberts, Jillian F. Banfield

**Affiliations:** Innovative Genomics Institute, University of California, Berkeley, CA, USA; Earth and Planetary Science, University of California, Berkeley, CA, USA; Bioengineering, University of California, Berkeley, CA, USA; Environmental Science, Policy and Management, University of California, Berkeley, CA, USA; Lawrence Berkeley National Laboratory, Berkeley, CA, USA; The University of Melbourne, VIC, AUS

## Abstract

Borgs are huge, linear extrachromosomal elements associated with anaerobic methane-oxidizing archaea. Striking features of Borg genomes are pervasive tandem direct repeat (TR) regions. Here, we present six new Borg genomes and investigate the characteristics of tandem repeats in all ten complete Borg genomes. We find that TR regions are rapidly evolving, recently formed, arise independently and are virtually absent in host *Methanoperedens* genomes. Flanking partial repeats and A-enriched character constrain the TR formation mechanism. TRs can be in intergenic regions, where they might serve as regulatory RNAs, or in open reading frames (ORFs). TRs in ORFs are under very strong selective pressure, leading to perfect amino acid TRs (aaTRs) that are commonly intrinsically disordered regions. Proteins with aaTRs are often extracellular or membrane proteins, and functionally similar or homologous proteins often have aaTRs composed of the same amino acids. We propose that Borg aaTR-proteins functionally diversify *Methanoperedens* and all TRs are crucial for specific Borg-host associations and possibly co-speciation.

## Introduction

Metagenomics has led to the increasing discovery of microbial extrachromosomal elements (ECEs) from environmental samples [1,2]. This bears great importance, as it allows us to better understand evolutionary processes and functional roles of ECEs in natural systems, and potentially use the ECEs, or elements of them, for genetic engineering. Using genome-resolved metagenomics we recently discovered Borgs, which are unusually large ECEs. Based on gene content and co-occurrence patterns, Borgs associate with several species of anaerobic methanotrophic (ANME) archaea of the *Methanoperedens* genus [3]. A search for other ECEs associated with these archaea also led to the recent discovery of large plasmids found in members of the same genus, yet a distinct clade of *Methanoperedens* species [4].

ANME perform anaerobic oxidation of methane (AOM) by using the reverse methanogenesis pathway [5]. One outstanding feature of Borgs is that they carry genes that encode proteins involved in key steps of their hosts’ metabolism. Remarkably, some Borgs encode the methyl-CoM reductase, all encode multiheme cytochromes (MHCs) that relay electrons onto extracellular terminal electron acceptors, and others encode nitrogenase used for nitrogen fixation [3]. The metabolic potential of different Borgs varies, suggesting diverse modes of interplay between Borgs and their hosts.

Borg genomes share a conserved genomic architecture that is very distinct from *Methanoperedens* chromosomes and plasmids of *Methanoperedens*. They are linear and large, with genome sizes for the first described examples ranging from 0.66 to 0.92 Mbp, thus exceeding known archaeal virus genome sizes by far. This places them in the range of giant and large eukaryotic double-stranded DNA viruses from the Nucleo-Cytoplasmic Large DNA Virus (NCLDV), with genome sizes that can exceed 2.5 Mbp [6]. Linearity also occurs in some virus genomes [7], and in eukaryotic chromosomes (*Saccharomyces*) as well as large linear plasmids of bacteria (*Streptomyces, Micrococcus luteus*) [8–10]. The Borg genomes reported to date have a large and a small replichore and both carry virtually all genes only on one strand. Borg genomes are terminated by long inverted repeats and perfect TRs are scattered throughout their entire genomes. These TR sequences follow a head-to-tail pattern, and occur in both intergenic and genic regions.

Here, we analyze Borg genomes to investigate the TR regions, focusing on those that comprise ≥ 50 nucleotides in length and include ≥ 3 repeat units. We found that DNA sequence assembly algorithms often collapse TR regions and that TRs frequently terminate contigs. This is not surprising, given that repeats in general are a well-known cause for errors in assemblies [7–10]. Thus, we augmented the four existing manually curated Borg genomes by manually curating six new Borg genomes, and twelve additional Borg contigs, to more completely uncover TRs. We found that TRs within ORFs caused perfect aaTRs within proteins. We then bioinformatically analyzed the aaTRs alone and in conjunction with the proteins they arise in. The high frequency, abundance and within population evolutionary dynamics of repetitive sequences suggest that they are fast evolving and have important biological functions.

## Results

### 2.1. Genome curation and TR analysis

#### 2.1.1. Curation of repetitive DNA sequences to complete Borg genomes

We reconstructed and manually curated six new Borg genomes to completion (see Methods; **Table 1**). Curation included correction of local errors where the automatically generated assemblies collapsed regions, or incorporated the wrong number of repeat units. These regions were identified visually based on elevated incidence of single nucleotide polymorphisms (SNPs) (**Figure 1A**) that clearly indicated that the region was misassembled (**Figure 1B**). All gaps created during the scaffolding step of the assembly were filled and genome fragments extended by making use of unplaced or misplaced paired reads (**Movie S1**). In some cases, the extending sequences were used to identify missing genome fragments that were then curated into the final assembly. In a few instances, TR regions exceeded the sequencing insert length so the real number of TR units could not be precisely determined. Resolving the *de novo* assembly errors unmasked TR regions in the genomes and revealed the aaTRs that these introduce into protein sequences.

**Table 1.** Tandem repeat (TR) statistics across 10 completed Borg genomes.

| Borg genome | nucleotide statistics |  |  |  |  |  | proteins statistics |  |  |  | TR<sub>orf</sub> /<br>TR<sub>intergenic</sub><br>(%) | TR<sub>intergenic</sub><br>div 3 (%) |
| --- | --- | --- | --- | --- | --- | --- | --- | --- | --- | --- | --- | --- |
|  | genome<br>size (bp) | # TR<br>regions | TR length<br>(bp) | TR /<br>genome (%) | # repeat<br>units | repeat per<br>bp | # proteins | # TR in<br>ORF | # aaTR proteins | aaTR proteins<br>/ proteome (%) |  |  |
| Black | 901883 | 56 | 6128 | 0.68 | 289 | 0.0003204 | 1292 | 23 | 26 | 2.01 | 41 | 21 |
| Brown | 937932 | 50 | 5913 | 0.63 | 259 | 0.0002761 | 1321 | 23 | 21 | 1.59 | 46 | 15 |
| Green | 1094519 | 76 | 10223 | 0.93 | 379 | 0.0003463 | 1517 | 30 | 26 | 1.71 | 39 | 15 |
| Lilac Borg | 661708 | 61 | 6733 | 1.02 | 299 | 0.0004519 | 825 | 28 | 22 | 2.67 | 46 | 33 |
| Orange | 974068 | 53 | 5675 | 0.58 | 246 | 0.0002525 | 1403 | 20 | 17 | 1.21 | 38 | 21 |
| Ochre | 725447 | 23 | 2108 | 0.29 | 92 | 0.0001268 | 1170 | 11 | 10 | 0.85 | 48 | 25 |
| Rose | 623782 | 20 | 2132 | 0.34 | 98 | 0.0001571 | 937 | 12 | 11 | 1.17 | 60 | 38 |
| Red | 685823 | 23 | 3181 | 0.46 | 173 | 0.0002523 | 1034 | 12 | 9 | 0.87 | 52 | 18 |
| Sky | 763094 | 32 | 3889 | 0.51 | 206 | 0.00027 | 1138 | 11 | 11 | 0.97 | 34 | 14 |
| Purple | 918293 | 66 | 6701 | 0.73 | 326 | 0.000355 | 1321 | 23 | 23 | 1.74 | 35 | 28 |
| average | 828655 | 46 | 5268 | 0.62 | 237 | 0.0002808 | 1196 | 19 | 17.6 | 1.48 | 44 | 23 |
| sum |  |  |  |  |  |  | 11958 |  | 176 |  |  |  |

**Figure 1.**
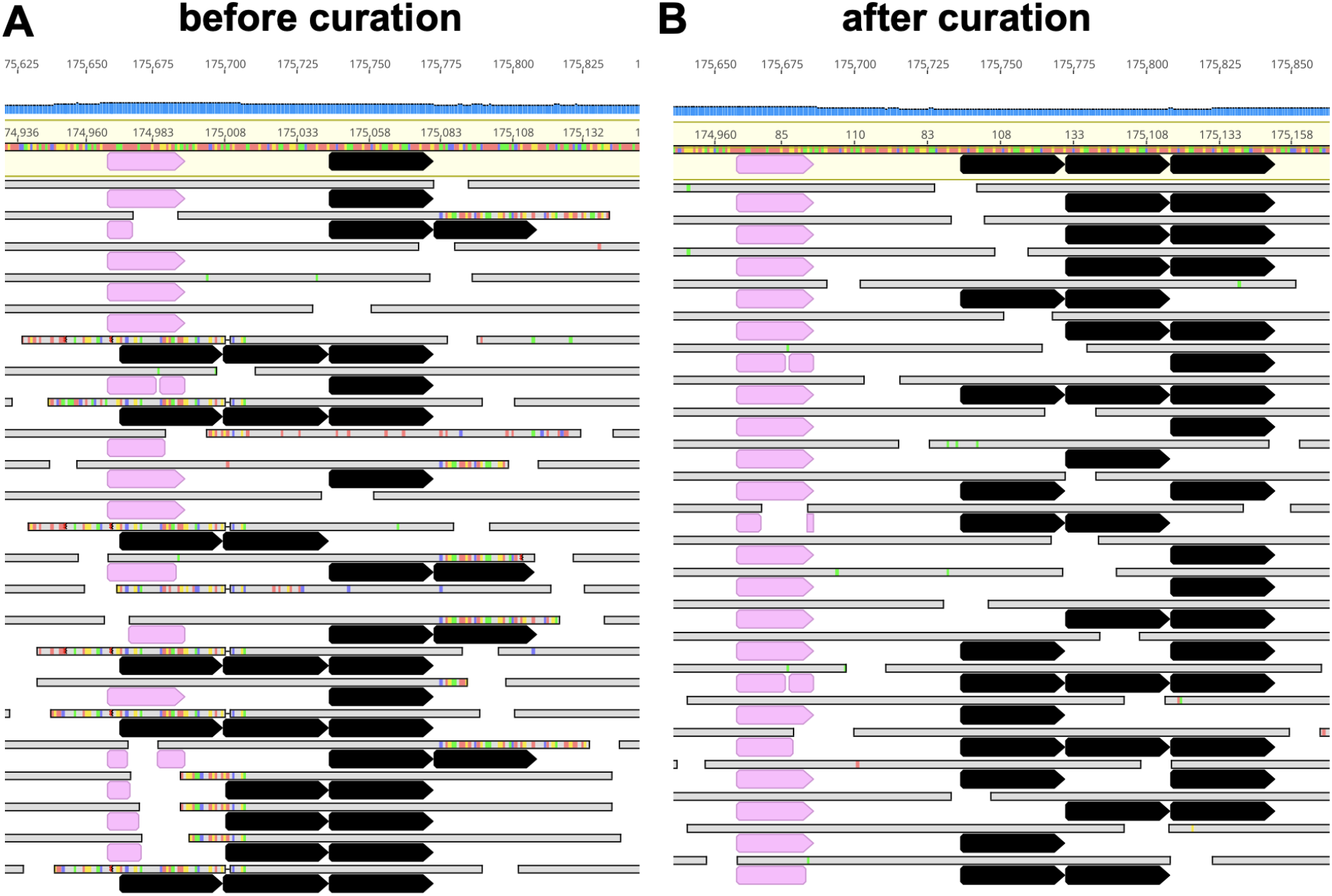
Reads mapped to a *de novo* assembly before and after curation. **A**. Large numbers of SNPs (marked as colored bars on reads shown in gray) indicate a local assembly error associated with a collapsed repeat region, resulting in a consensus sequence (top) with one repeat unit (black). **B**. The consensus sequence has three repeat units after curation. The pink segments mark a unique sequence that was used to place misplaced reads correctly.

Rose Borg possesses the smallest (623,782 kbp) and Green Borg possesses the largest genome (1,094,519 kbp). As expected based on the four previously reported genomes, five of the new genomes are linear, terminated by inverted repeats. Based on GC skew analysis, replication of Borg DNA is initiated at the chromosome ends (**Figure 2, Figure S1**). The Red Borg genome contains a repeated sequence that prevented identification of a unique genome assembly path. The variant that generated the expected pattern of GC skew as for the other Borgs was chosen, completing the final set of ten Borg genomes. All genomes but the Green Borg genome are composed of a large and a small replichore; the Green Borg genome has a slightly more complex organization. Each replichore carries essentially all genes only on one strand. Consequently there are no apparent transcriptional operons. This was mirrored by a low frequency of transcriptional regulators in Borg genomes. Specifically we only found 0.35 transcriptional regulators per 100 kbp Borg genome, whereas near complete *Methanoperedens* genomes encoded 5.9 in 100 kbp (1 in Rose, 2 in Purple, 3 in Brown, Lilac, Sky, Green, 5 in Purple, 6 in Orange and none in Ochre and Red Borg versus 53, 53 and 62 in three near complete *Methanoperedens* genomes SRVP18_hole-7m-from-trench_1_20cm__Methanoperedens-related_44_31,RifSed_csp 2_16ft_3_Methanoperedens_45_12,RifSed_csp1_19ft_2_Methanoperedens_44_10; 2.77 - 2.90 Mbp). We also note that none of the Borgs encode a DNA-dependent RNA polymerase or a TATA-box binding protein.

**Figure 2.**
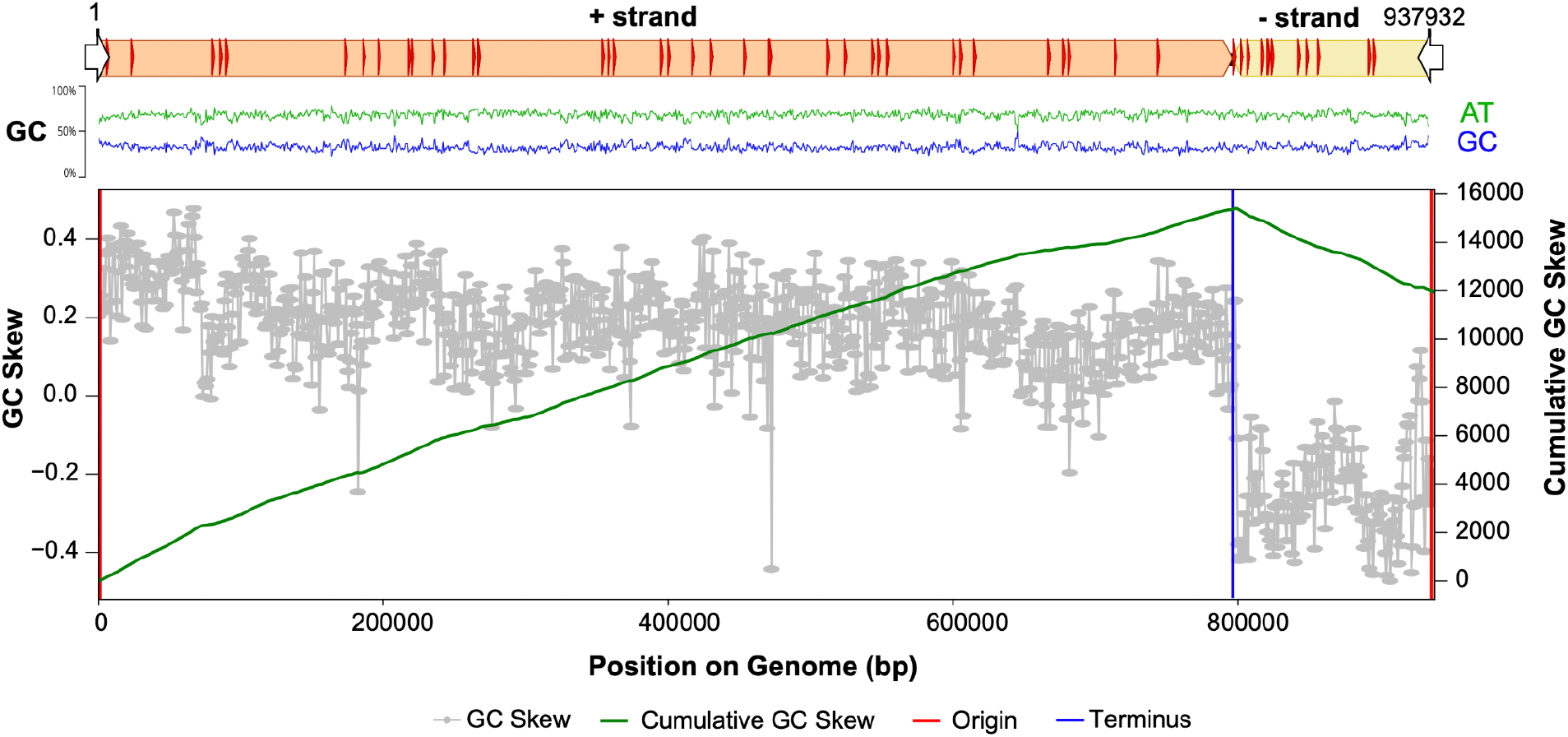
Borg genomes are composed of a large and a small replichore and their replication is initiated at the termini. Shown is the genome architecture of Brown Borg, including terminal inverted repeats (white arrows), TR regions (red arrows), and GC composition. GC skew and cumulative GC skew show that the genome is replicated from the terminal inverted repeats (origin) to the terminus.

The previously reported four completed Borg genomes and six new completed Borg genomes were used to evaluate the distribution and features of perfect TRs using a stringent threshold that allowed no mismatch (≥ 50 nt region and ≥ 3 TR units). However, during the manual curation we noticed that many regions had mapped sequencing reads with slight differences in the unit composition, usually a single nucleotide change. In the example of a repeat region in a MHC gene, there is a G → A substitution SNP that is synonymous on the amino acid level (**Figure S2**). Interestingly, the repeat variants may not occur sequentially and the mixture of variants can be different for a single Borg population.

#### 2.1.2. Regions with TRs are fast evolving

The number of TR units in a region sometimes clearly differed within a single Borg population (**Figure 3A, 3B**). This implies that the Borg TR regions are fast evolving, much like CRISPR repeat-spacer inventories. Moreover, the repeat loci were rarely conserved in otherwise alignable regions of the most closely related Borgs (and were absent from homologous proteins from organisms). A rare case where regions are similar occurs in Black and Brown Borgs, which are closely related based on genome alignments (**Figure 3C**). Close inspection of a ~7 kbp region in the aligned genomes revealed four different kinds of TRs. TR-1 is intergenic, is composed of a 20 nt unit repeated six times, followed by a nearly perfect additional unit of 21 nt, then another perfect unit. This intergenic TR is absent in Brown Borg. TR-2 is in an ORF and comprises 18 nt units that are identical in Black and Brown Borgs, where they occur seven and eight times, respectively. TR-3 is also in an ORF and comprises 21 nt units that occur consecutively six times in Black Borg. Brown Borg has two identical units in the same ORF, followed by a sequence that differs by one SNP, then another identical unit. TR-4 is also in an ORF and comprises 36 nt units that occur four times in Brown Borg, but these are absent in the same ORF from Black Borg. Interestingly, the nucleotide sequence from Brown Borg in the vicinity of TR-4 has high nucleotide-level similarity but no TRs.

**Figure 3.**
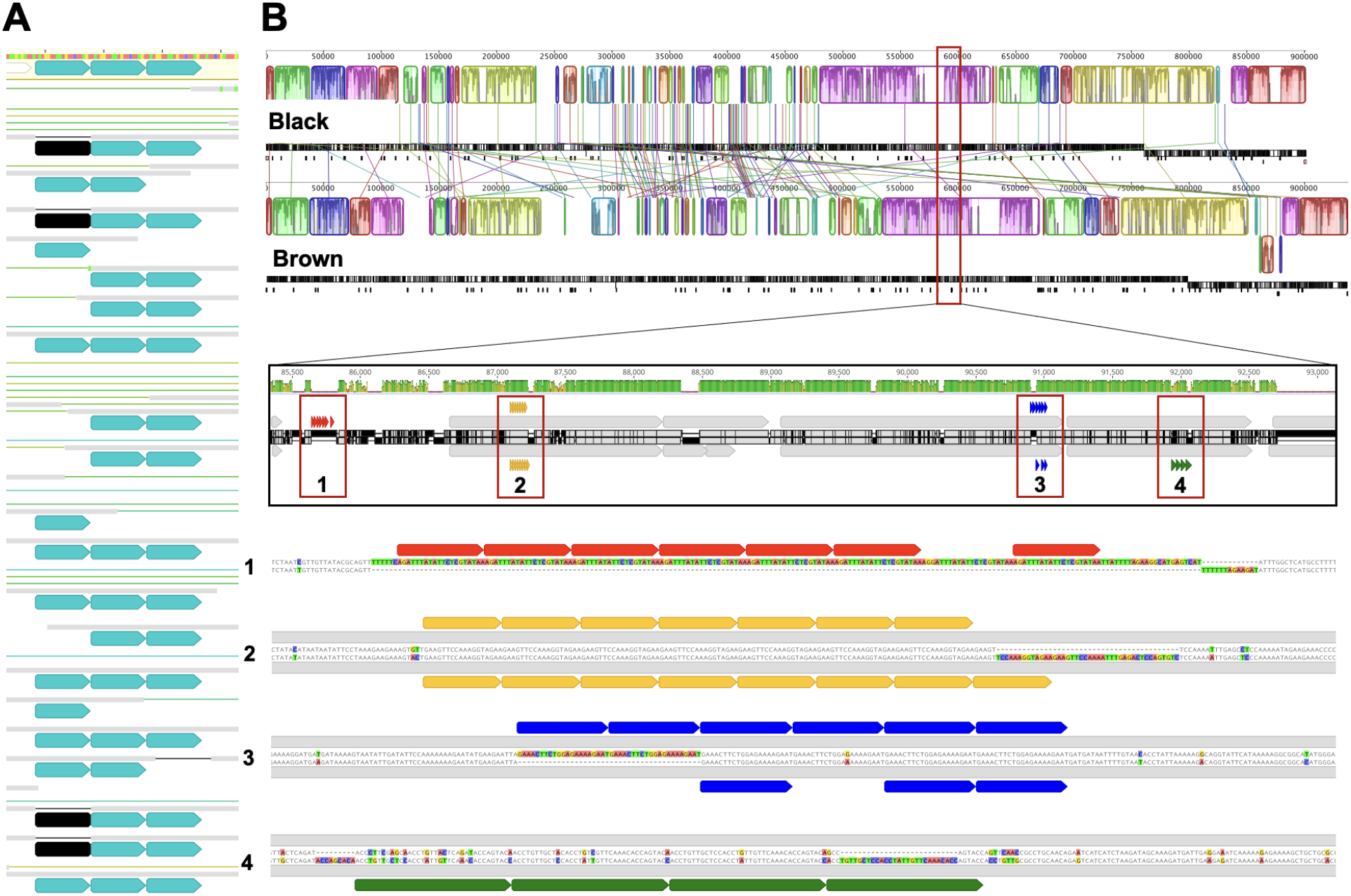
Subpopulation variability of TR unit numbers and population differences in TR regions. **A**. Four reads show different TR unit numbers than the consensus sequence (top) within a single Borg population. TR units in aqua, missing TR units in black. **B**. Genome alignment of closely related Black Borg and Brown Borg. A 7 kb region reveals four different TR instances (small red boxes) in an otherwise homologous region.

#### 2.1.3. TRs are flanked by partial repeats and are enriched in adenosine

To constrain the mechanism behind tandem repeat formation in Borg genomes, we assessed characteristics of the TR sequences and their flanking DNA regions. Repetition of sequences with variation in GC versus AT content generally introduces GC/AT-symmetry in regions containing TR units (**Figure 4**). Offset of the symmetric units and the TRs depends on the exact choice of the repeat unit. The TR regions can be preceded by sequences that are identical to the end of the TR unit and/or followed by sequences that are identical to the start of the TR unit (**Figure 4A, B**). These flanking partial repeats are often also adjacent to TR regions that contain only two repeats (and were excluded from statistics). Often the middle of the TR unit is not in either of the 5’ or 3’ partial repeats. When there are different choices for the repeat unit there are slightly different partial repeats that flank the TR sequences (**Figure 4C**).

**Figure 4.**
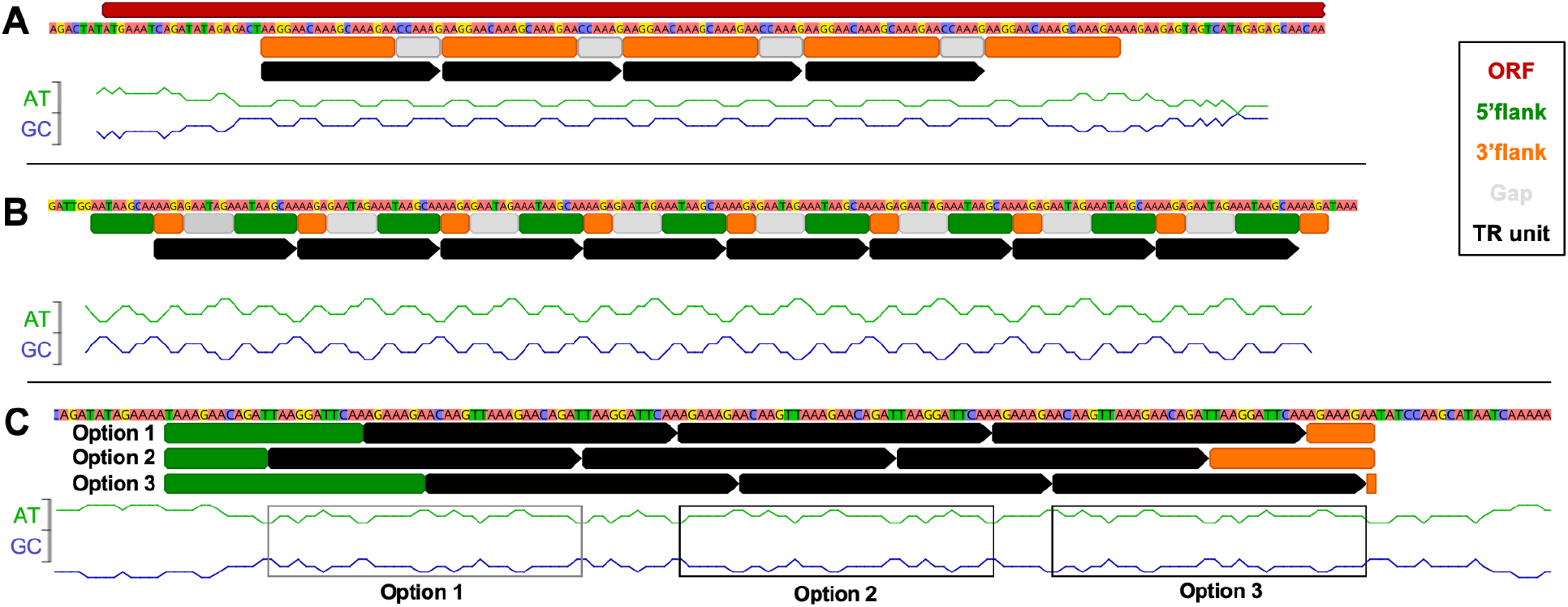
Tandem repeats are often flanked by partial repeat sequences. **A**. TR regions can occur within ORFs. **B**. TRs can be in intergenic regions (or the region can start within the end of an ORF). The TR regions are often flanked by partial repeats. **C**. Different options for the start of a TR region define different flanking partial repeats.

The nucleotide composition of the ORF-carrying strands on both replichores was similar, with nucleotide frequencies of A>T>G>C (38.4%, 28.8%, 20.3%, 12.5%). The nucleotide composition of all TR units compared to this overall composition showed a strong bias towards A (48.4%) while all other nucleotides were depleted (T 23.7%, G 17.1%, C 10.8%). This was less pronounced in TRs within ORFs (A 46.3%) and these TRs were also enriched in C (15.5%), whereas T and G were depleted (18.6%, 19.6%) (**Table S1**). The fact that this compositional bias is observed in both genes, intergenic regions and in flanking partial repeats suggests that the A enrichment is important for the process that forms the repeats. Notably, we did not find a uniform secondary structure of tandem repeats, since some formed predicted loops and others formed predicted hairpins.

We searched for polymerases in Borg genomes, given their potential relevance for repeat formation. We found that all Borgs encode at least one DNA polymerase and phylogenetic placement within reference sequences [11] revealed that there are two types of PolBs encoded in the Borgs: the B2 clade and the B9 clade (**Figure S3**). The DNA PolB9 encompasses a predicted 3’ to 5’ exonuclease, and was found in each complete Borg genome. The amino acid sequences are very similar, suggesting a high mode of conservation for these particular proteins.

#### 2.1.4. TRs are rare in *Methanoperedens* and introduce aaTRs in Borg ORFs

To shed light on the role and function of the tandem repeats, we surveyed their occurrence across all ten complete Borg genomes. We found 460 regions that make up 0.62% of the average Borg genome (**Table 1**). Draft *Methanoperedens* metagenomes on the other hand only have 1-4 TR regions, making up ≤ 0.01% of the metagenomes (0.0099% in SRVP18_hole-7m-from-trench_1_20cm__Methanoperedens-related_44_31, 0.0021% RifSed_csp2_16ft_3_Methanoperedens_45_12, 0.0018% in RifSed_csp1_19ft_2_Methanoperedens_44_10), suggesting that TR formation is highly genome specific. Approximately half (43-65%) of the Borg TRs were located within ORFs. All TRs within ORFs had unit lengths that are divisible by three, so these repeats do not disrupt reading frames. They result in amino acid tandem repeats that we refer to as aaTRs (and aaTR-proteins). Only 14-38% of intergenic TR unit lengths were divisible by three (**Figure S4**). Sometimes several different TRs occurred within the same ORF, so the dataset comprised 214 aaTRs in 178 individual proteins from 10 Borg genomes. Fifteen aaTR-proteins were not perfect TRs on the nucleotide level, yet almost all were imperfect due to the occurrence of single SNPs. The TR nucleotide sequences are virtually all unique within each Borg genome and are very rarely shared between Borg genomes. Upon inspection of aaTR-proteins, we found a different codon usage of genes with TRs relative to all Borg genes. As expected based on the relatively A-rich composition of repeat regions, the aaTR-bearing ORFs favored incorporation of codons that were often higher in A than T, G, C (**Figure S5**).

### 2.2. Biophysical properties of aaTRs

#### 2.2.1. Amino acid frequency in aaTRs

We closely examined the predicted biophysical and biochemical properties of the single repeat units from all ten Borg genomes and 37 additional proteins from curated Borg contigs. This final dataset comprised 215 Borg aaTR-proteins and 306 repeat regions (**Table S2, S3**). Proline, glutamate, glutamine and lysine were particularly enriched across all Borgs, while cysteine, phenylalanine and tryptophan were almost absent (**Figure 5A**). We noticed that disorder-promoting amino acids were enriched in the aaTRs and order-promoting amino acids were depleted (**Figure S6**).

**Figure 5.**
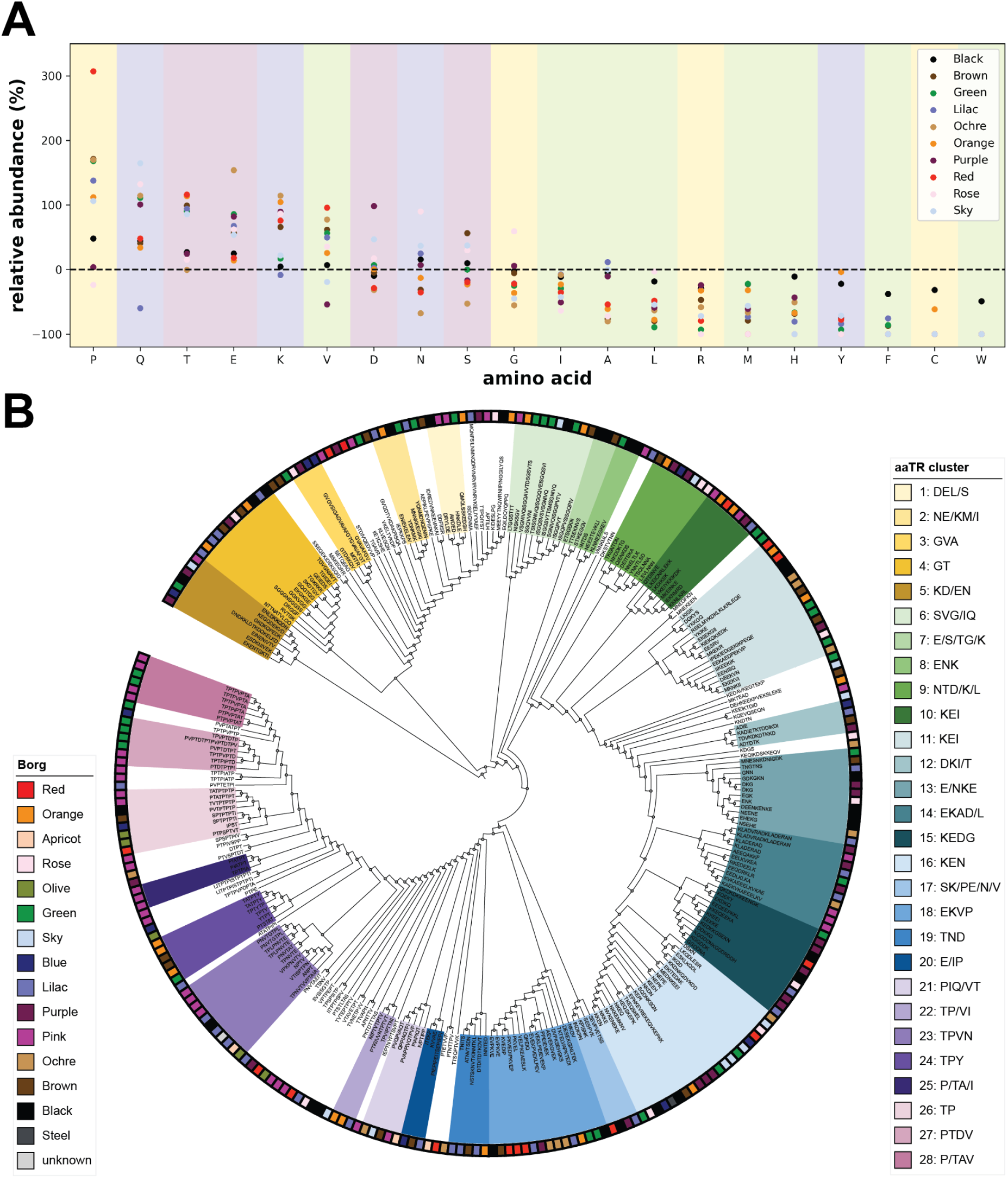
Relative amino acid abundance in aaTRs and hierarchical clustering of triple aaTR units. **A**. Relative aa abundance in aaTRs compared to all proteins. Background colors indicate the aa properties: charged (lilac), polar uncharged (magenta), special cases (yellow) and hydrophobic (green). **B**. Repeat units were hierarchically clustered based on triple aaTR units. Clusters with ≥ 3 aaTR members were named after descending aa abundance (≥10%). The color strip shows in which Borg genome each aaTR is encoded.

Hierarchical clustering of triple aaTR units revealed 28 aaTR clusters, each cluster composed of aaTRs with very similar amino acid composition and frequency (**Figure 5B**). Many aaTR units possessed nearly equal numbers of the charged amino acids K and E (polyampholyte repeat cluster), others were enriched in P/T/S, which are potential sites of post-translational modification. From this we conclude that there are distinct and related groups of aaTRs. The related groups share common properties and may fulfill similar functions.

#### 2.2.2. aaTRs are enriched in extracellular proteins and most aaTRs are intrinsically disordered

Strikingly, 32.1% (69/215) of aaTR-proteins have extracellular regions and 30.2% have transmembrane helices (TMHs), whereas only 18.6% (2,230/11,995) and 16.7% of all Borg proteins have these features. Consistently, aaTR-proteins are highly enriched in signal peptides (16.7% of aaTR-proteins compared to 4.0% of all Borg proteins).

To further investigate the observation that aaTRs particularly often contain disorder-promoting amino acids, we assessed the prominence of intrinsically disordered regions (IDRs) in aaTR-proteins versus all Borg proteins. IDRs are polypeptide segments that are characterized by a lack of a well-defined 3-D structure [12]. Remarkably, 62.8% aaTR-proteins contained at least one IDR, whereas only 5.6% of all Borg proteins contained an IDR. The IDRs almost always corresponded to the TR region, and *Methanoperedens* homologues did not have IDRs (**Figure 6A**). Thus we hypothesize that the TRs in ORFs mostly lead to the creation of new intrinsically disordered regions in existing proteins.

**Figure 6.**
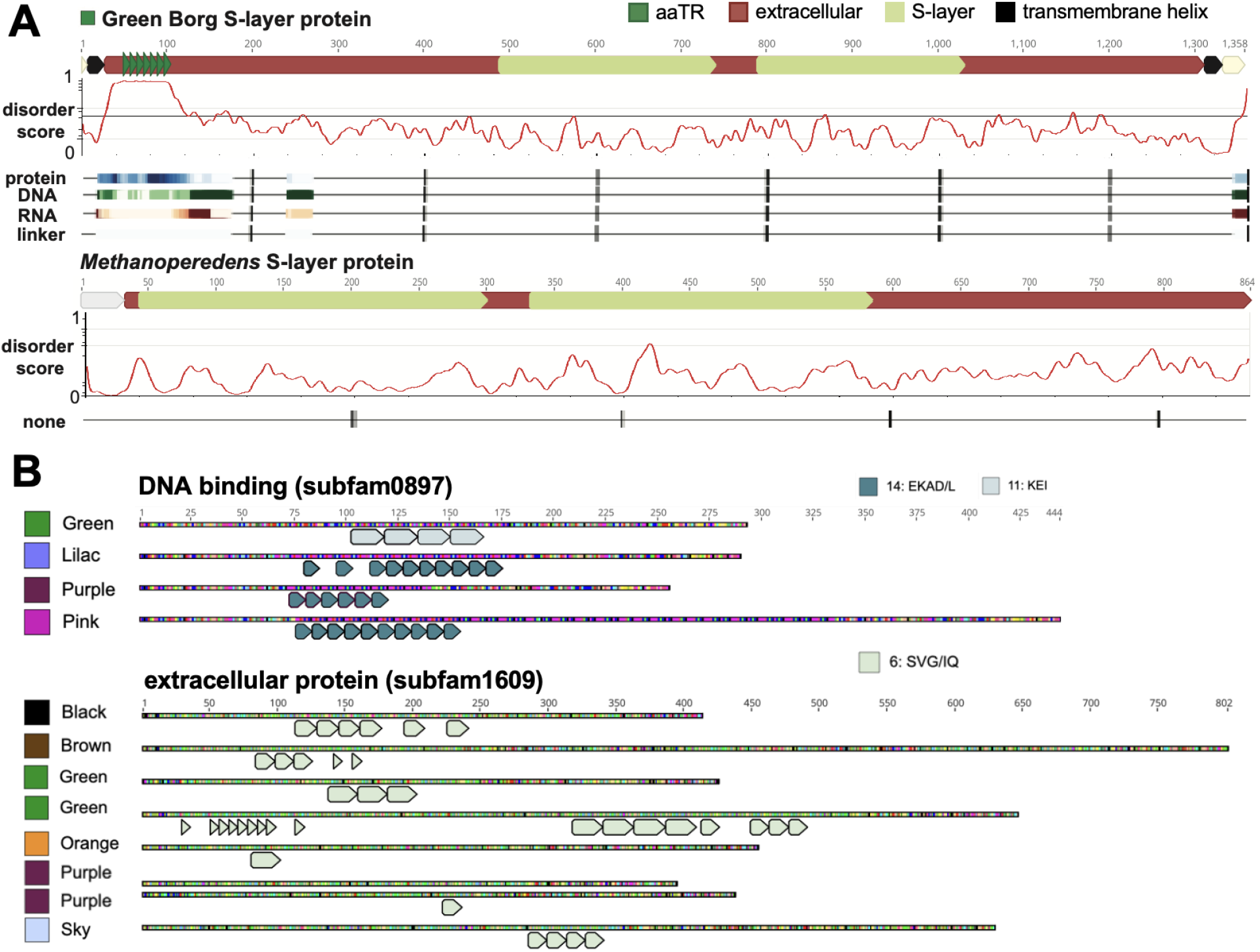
aaTRs introduce intrinsic disorder, are predicted binding sites and have the same aa composition in homologous Borg proteins. **A**. Domain architecture of S-layer proteins from Green Borg and a *Methanoperedens* homologue (WP_096205607.1). The aaTR is an IDR and a predicted binding site for protein, DNA, RNA, or a linker region. This region is absent in the *Methanoperedens* homologue. Intrinsic disorder score, binding and linker propensity were calculated with IUPred2A and flDPnn. **B**. All or most members of two protein subfamilies have aaTRs. The aaTRs within a protein subfamily are from one or two related aaTR clusters.

Forty proteins possessed multiple repeat regions ranging from two to five aaTR regions (**Table S4**) and the aaTRs were often located at the N- or C-termini. Some of these regions fall into the same or a similar aaTR cluster, whereas others are distinct and often located in a completely different region of the polypeptide chain. In 22 cases, the aaTRs make up the majority (50 - 96%) of the predicted proteins and many of these are small proteins (14 proteins are < 100 aa) (**Table S5**). The repeat units of these proteins are diverse, but clusters enriched in K, E and I (cluster 10 and 11) were particularly frequent (6/18). Although it is possible that these are wrongly predicted genes in intergenic regions, they may also be *de novo* repeat peptides. Their existence as real proteins is supported by the observation that they possess a start codon (or alternative start codon), the TR units are divisible by 3 and thus introduce aaTRs, and by the fact that four have signal peptides and four have functional annotations (linked to apolipoprotein, proline-rich domains and a glycoprotein domain).

### 2.3. Functions of aaTR-proteins

#### 2.3.1. Similar aaTRs fulfill similar predicted functions and functionally related proteins have similar aaTRs

To investigate which functions aaTR-proteins have, and if functionally related proteins possess similar aaTRs, we performed protein family clustering of 11,995 Borg proteins. 85% of Borg proteins were grouped into 1,890 subfamilies and 80% (172/215) of the aaTR-proteins clustered into 112 subfamilies (**Table S1**). The functional landscape of the subfamilies ranged from carbon metabolism (18 proteins), cell and protein architecture and scaffolding (9), nucleotide processing (15), transcription-associated proteins (8), redox (8), signaling (11), transport (3), stress response (2), to protein processing (1) (**Table 2**).

**Table 2.**
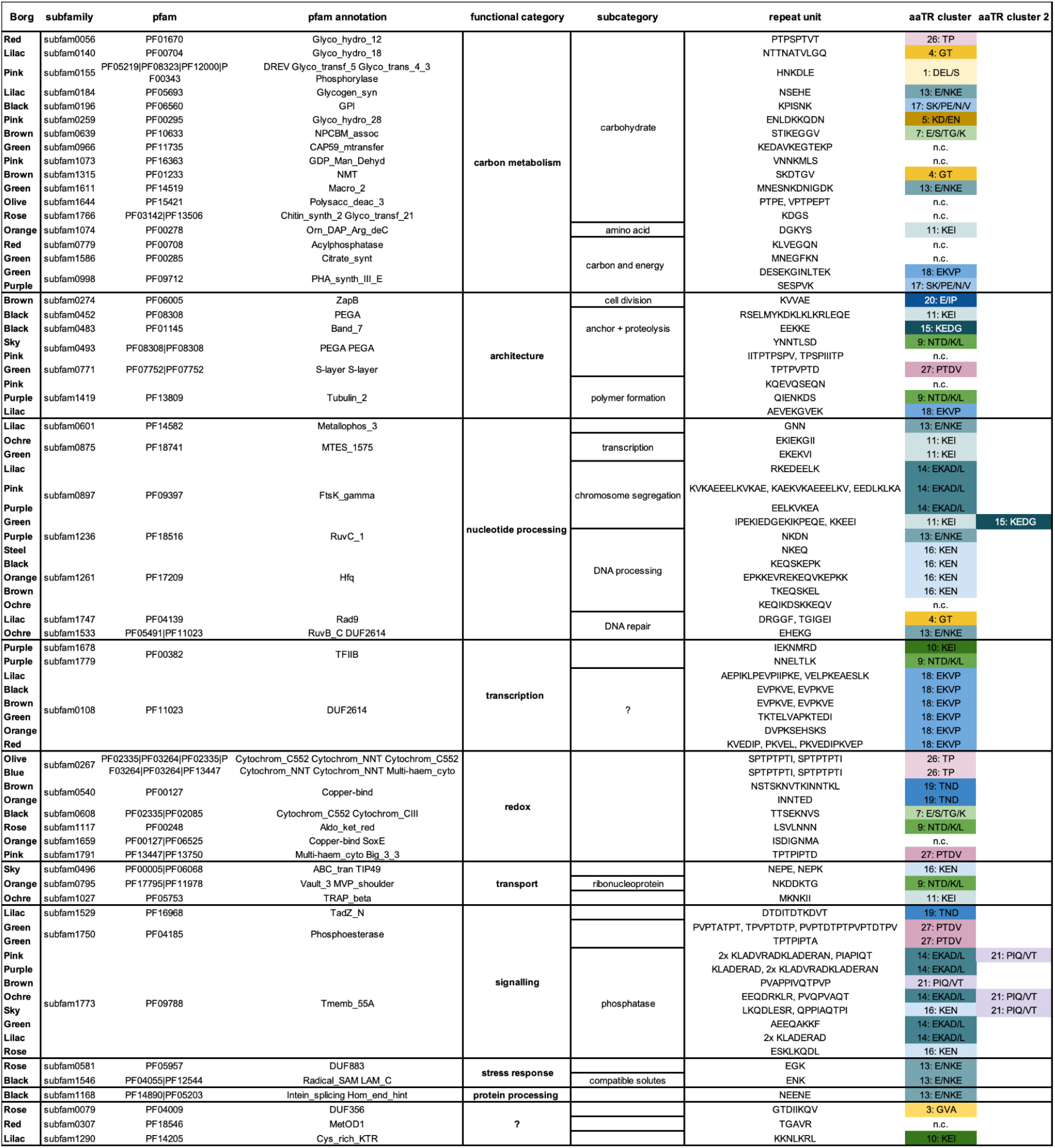
Functional annotations for subfamilies with aaTR-proteins. Proteins were manually placed into functional categories based on the pfam annotation of the subfamily. Single aaTR units are listed as well as in which aaTR cluster they fall. Some aaTRs did not fall into clusters (n.c., no cluster), and some proteins have two or more aaTRs that fall into one or two clusters.

Screening the aaTR-proteins with the eukaryotic linear motif (ELM) resource for functional sites in proteins (http://elm.eu.org/) unmasked that the aaTRs within them have similarity to a plethora of different short linear motifs (SLiMs). SLiMs are composed of three to ten consecutive amino acids [13], which are used by eukaryotic cells as cleavage sites, degradation sites, docking sites, ligand binding sites, post-translational modification sites and targeting sites [14]. We found that aaTRs from the same aaTR cluster or related aaTR clusters often had matching functional predictions (see examples in **Table S3**).

Fifteen protein subfamilies were enriched in aaTR-proteins, which included subfamilies functionally annotated as transmembrane phosphatases, ribonucleoproteins, phosphoesterases, zinc-ribbon proteins and DNA-binding proteins; the other subfamilies had no functional annotations (**Table S2**). Members of the same protein subfamily often had similar aaTRs. For example, a subfamily of DNA binding proteins (subfam0897) only comprises aaTR-bearing members, all of which have unique aaTRs, yet of the same or related repeat clusters (**Figure 6B**). These aaTRs form a predicted coil structure or an IDR and resemble SLiMs that play a role in localization, protein stability or phosphorylation-dependent signaling. Similarly, subfam1609 comprises eight members, five of which are aaTR-proteins and despite all being only distant homologues (highest aa identity is 67%), the aaTRs all belong to the same aaTR cluster (SVG/IQ) which corresponds to predicted modification (phosphorylation) and ligand-binding sites (**Figure 6B, Table S3**).

#### 2.3.2. aaTR-proteins responsible for cell integrity/stability and surface decoration

Several functionally related, but phylogenetically unrelated proteins that are responsible for cell wall architecture (PEGA and S-layer) and decoration (glycosyltransferases and glycosyl hydrolases) have aaTRs. The aaTRs of the PEGA proteins resemble predicted ligand binding, docking and modification (phosphorylation) sites and the aaTRs of the S-layer resembles proteolysis-initiating degrons [15]. The aaTRs of glycosyl hydrolases resemble modification and ligand binding sites and the aaTRs of glycosyl transferases resemble degrons (**Table S3**). Some Borgs also possess an aaTR in tubulins, which are proteins required for cell division [16]. These aaTRs resemble SLiMs that are potential docking and cleavage sites and/or initiate proteasomal degradation. Importantly, these SLiM-bearing aaTRs are absent in non-Borg homologues.

### 2.4. Case studies of aaTR-proteins: ribonucleoproteins, MHCs and a conserved aaTR hotspot

To further investigate proteins with the same function that evolve similar but not identical aaTRs, we performed in-depth analyses of two distinct types of proteins with a known function and a conserved region across Borgs comprising multiple aaTR-proteins.

#### 2.4.1. Ribonucleoproteins

Most Borgs encode Sm ribonucleoproteins, which are archaeal homologues of bacterial Hfq and eukaryotic Sm/Sm-like (Lsm) proteins. These proteins are implicated in versatile functions such as RNA-processing and stability and the protein monomers form a stable hexameric or heptameric ring-shaped particle [17,18]. We found 19 Borg Sm ribonucleoproteins (**Table S6**), five possess aaTRs and one additional sequence from Grey Borg has a near-perfect aaTR. The aaTRs are always located at the N-terminus, preserving the conserved Sm1 and Sm2 RNA-binding motifs (**Figure S7A**). The aaTR units contain 4, 8, 12 or 17 amino acids, have the same aa character, and are predicted SLiMs that facilitate docking or resemble degrons (**Figure S7B, Table S6**). An initial homology-based structural search of the aaTR-Sm from Black Borg identified the bacterial Hfq of *Pseudomonas aeruginosa* (PDB: 4MML, aa identity: 26%) that forms a homohexameric ring structure. Due to no alignable template in the database for the aaTR, the model failed to predict a structure of this N-terminal region. Remodeling with AlphaFold2 predicted the structure of the Sm core that formed a hexameric ring with N-terminal repeat regions that extend as long loops on the distal side of the protein (**Figure 7**). Modeling of the other four aaTR-Sm proteins also showed N-terminal unstructured extensions corresponding to the aaTRs (**Figure S7C**).

**Figure 7.**
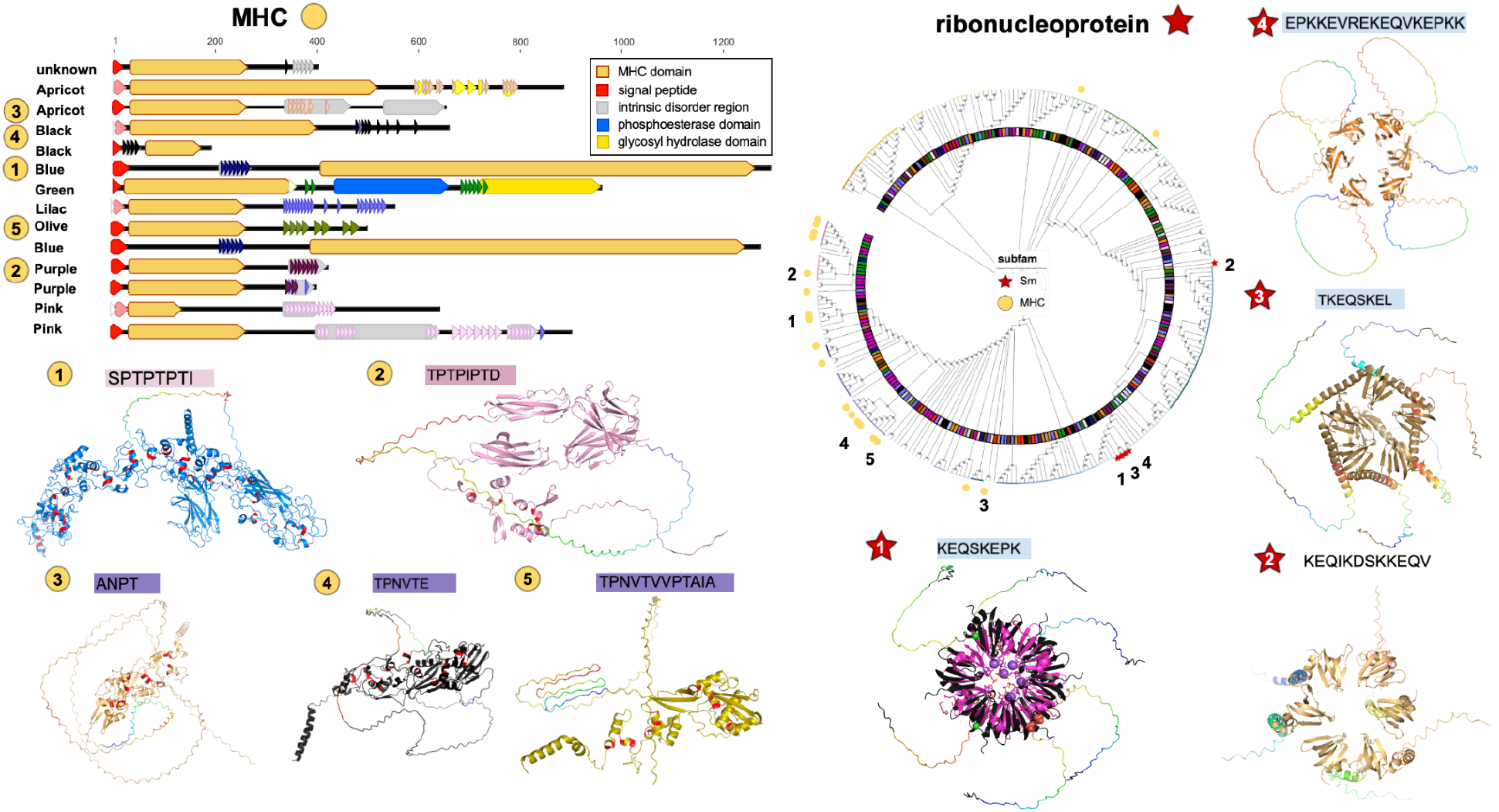
Domain architecture and predicted structure of aaTR-bearing MHCs and ribonucleoproteins. aaTR units are depicted as colored arrows in the domain topology panel. Predicted structure of select aaTR-MHCs with CxxCH and aaTR regions highlighted in red and rainbow. Sm ribonucleoprotein from Black Borg (1, black) was superimposed on Sm from *M. jannaschii* (PDB: 4×9D with UMP RNA + ions, magenta). aaTR regions in Sm ribonucleoproteins from Ochre (2), Brown (3) and Orange (4) Borg are highlighted in rainbow.

#### 2.4.2. Extracellular electron transferring MHCs

All Borgs encode MHCs, which are particularly important and abundant in *Methanoperedens* as they mediate the final electron transfer from methane metabolism to an external electron acceptor [19]. We found 14 MHCs with aaTRs in Borg genomes. All of these are predicted to be located extracellularly, some possess a membrane anchor, and they range from 20.5 kDa (190 aa) to 137.5 kDa (1,293 aa) and contain up to 30 heme binding sites (**Figure 7**). The aaTRs are located at different sites in the polypeptides, but nevertheless belong to identical or similar repeat clusters that are consistently enriched in T and P and resemble ligand binding, docking and modification sites (**Table S7**). The aaTRs are mostly IDRs which are predicted to form unstructured extensions that protrude from the folded protein core.

#### 2.4.3. A conserved aaTR hotspot

There is a region in all completed Borg genomes that is a hotspot for TRs with up to five aaTR-bearing proteins. It includes a gene encoding subfam1773, which we refer to as cell envelope integrity protein TolA. Best blastp and structural hits are TolA from, e.g., the pathogenic bacterium *Leptospira sarikeiensis* (48% aa identity) and YgfJ from the pathogen *Salmonella typhimurium*. Eight TolA proteins have unique aaTRs, and some carry additional aaTR units shared by other Borgs (e.g., Ochre Borg repeat units are found in Rose Borg and Purple Borg; **Figure 8A, C**). These aaTRs are predominantly predicted degradation, docking and ligand binding sites (**Table S7**). The regions encode two other conserved protein subfamilies with aaTR-bearing members: subfam0649 lacking functional annotation and a subfamily of zinc-ribbon proteins (subfam0108), which usually form interaction modules with nucleic acids, proteins or metabolites (**Figure 8B**). The subfam0108 proteins possess seven distinct aaTR units, six of which fall into the same repeat cluster (**Figure 8C**). Additional aaTR-bearing proteins in the same context are a glycogen synthase/glycosyl transferase (subfam0184) in Black Borg and Green Borg and proteins of unknown function (subfam1382) in Lilac Borg and Orange Borg, and a hypothetical protein in Green Borg.

**Figure 8.**
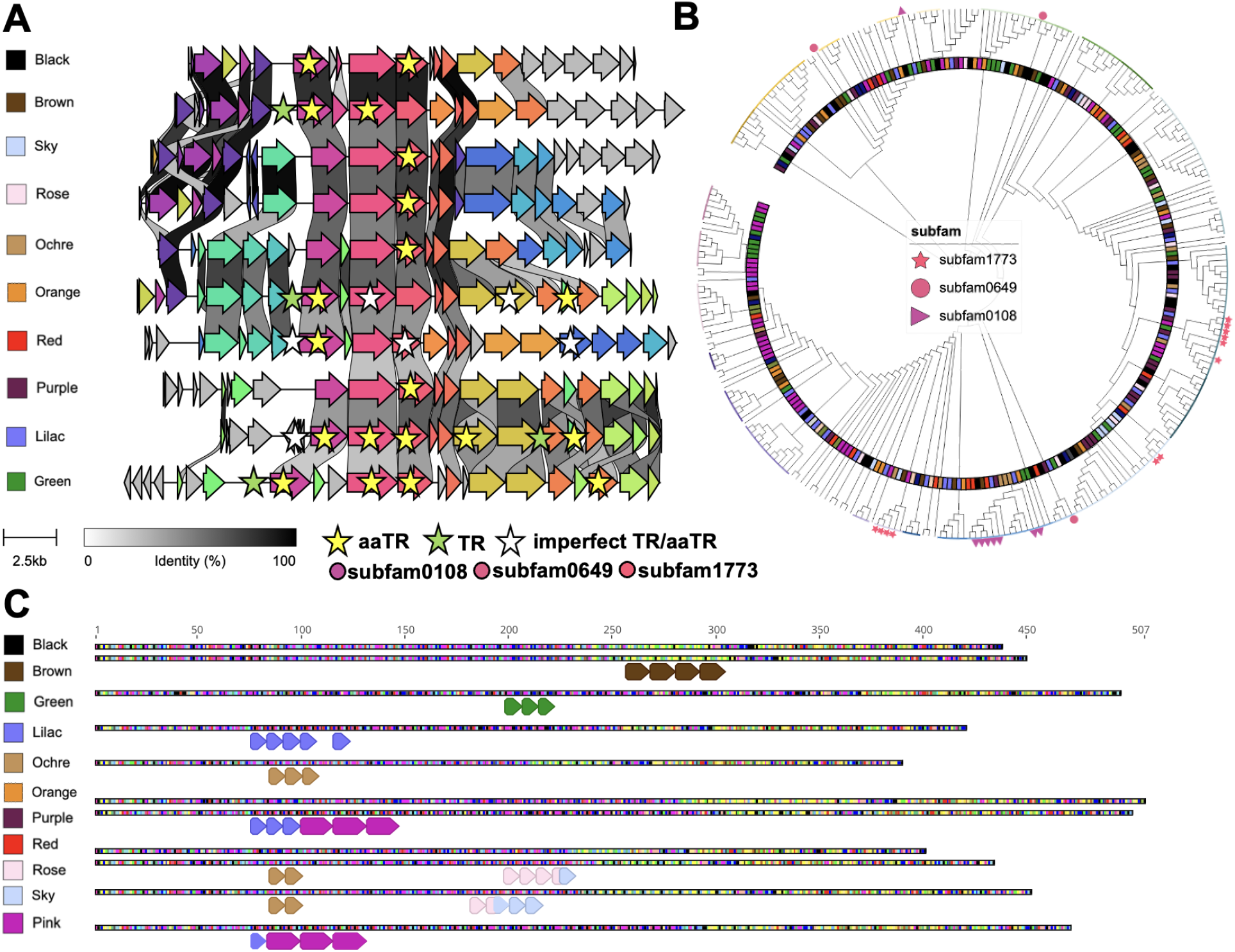
A conserved genetic region is a hotspot for tandem repeats. **A**. Amino acid alignment of a conserved genetic region that encodes TolA (subfam1773). **B**. aaTRs from subfam0108, subfam0649 and subfam1773 cluster together. **C**. Eight out of eleven Borg TolA homologues have aaTRs.

## Discussion

Tandem nucleotide repeats in Borg genomes likely are formed and evolve independently, as evidenced by the essentially unique sequences of tandem repeat units in each region within and among Borg genomes. They evolve rapidly, based on differences in the alignable regions of the most closely related Borgs (e.g., Black and Brown Borgs) and variability in repeat numbers within a single Borg population. This parallels the rapid evolution of eukaryotic variable number tandem repeats (VNTRs). VNTRs are widespread and linked to neuropathological disorders (caused by the accumulation of short tandem repeats) [20], gene silencing [21], and rapid morphological variation [22]. In bacterial genomes, highly repeated sequences are less common, and TR variations are linked to immune evasion, cell-pathogen specificity (definition of which cells/tissues/host pathogens/viruses can infect) and stress tolerance [23,24]. They are relatively common in a few archaeal genomes, but the functions there remain uncertain.

Previously proposed mechanisms for the expansion and retraction of VNTRs implicate various DNA transactions, including replication, transcription, repair and recombination [25]. Replication slippage is one explanation for the origin and evolution of repeat loci [23]. This mechanism involves a pausing of the DNA polymerase and dissociation from the TR region, followed by DNA reannealing. Realignment of the newly synthesized strand can be out-of-register on the template strand, leading to propagation/retraction of the TRs. At this time, it is unclear whether the TRs are introduced by Borg or *Methanoperedens* machinery. If the former, it is notable that we found that all ten Borgs encode a highly conserved DNA PolB9, a clade of uncharacterized polymerases that have only been reported from metagenomic datasets [11]. Archaeal DNA polymerases B and D have been shown to slip during replication of tandem repeat ssDNA regions [26]. Thus the Borg PolB9 could be responsible for TR propagation, possibly triggered by non-canonical secondary structures formed by the tandem repeat DNA or its upstream region [26].

Given the shared nucleotide characteristics of genic and intergenic TR regions (perfection, A bias) we infer that they form by the same mechanism. Even if TRs are introduced perfectly, one would expect some of them to have accumulated mutations, unless perfection is strongly selected for and ensured through a repair mechanism, or unless they all formed very recently. The latter is consistent with other evidence that TR regions are fast evolving.

In many cases, the sequences that flank the TR regions are partial repeats. We suspect that these are seed sequences that were used to initiate TR formation. If these regions were the seeds that gave rise to the TRs, the fact that they are often only parts of the repeat raises a mystery regarding the origin of central regions of these TRs. It is possible that the sequences that served as seeds were subjected to elevated mutation rates, which is supported by the observation of high SNP levels flanking TR regions in the human genome [25,27].

CRISPR repeats suggest a possible alternative to the replication slippage explanation for the origin of TR regions. Like TR regions, CRISPR loci are fast evolving, in that case motivated by the need to counter rapid evolution of bacteriophages to outwit spacer-based immunity [27]. CRISPR repeats are introduced by an integrase that excises the previously added repeat, ligates it to a new spacer, and adds that unit to the expanding locus, filling gaps to recover a double stranded sequence [28]. If a Borg system is involved, the genomes encode many and various nucleases and recombinases, some of which may be responsible for repeat addition. Unfortunately, a complete *Methanoperedens* genome is not available yet, so we cannot confidently assess their capacities for repeat introduction.

As noted above, our analyses revealed a compositional bias in nucleotide tandem repeats towards A on the coding strand of each replichore, primarily at the expense of T. If this reflects mutation, the transversion (conversion of purine to pyrimidine) must happen at the biogenesis of the TR template, as the TRs are faithfully copy-pasted error-free. We tentatively support the alternative explanation that the A-rich nature of the seeds initiated TR formation. A small region that is A-rich, possibly functioning in a manner somewhat analogous to a protospacer adjacent motif (PAM) that binds the CRISPR system nuclease, may localize the machinery involved in repeat formation. We speculate that TR formation could be regulated by distinct methylation patterns of Borg DNA, possibly related to the A-rich character of the seed.

A key question relates to how stable coexistence of Borgs, with their huge genomes, likely in multi-copy [3], and the host *Methanoperedens* cells is achieved. Although it is possible that Borgs direct the functioning and evolution of the hosts upon which they fundamentally depend on for their existence, it is perhaps more likely that *Methanopredens* controls Borg activity. The mechanisms for regulation of Borg gene expression are unclear, given the absence of obvious operons and the conspicuous lack of transcriptional regulators. As we identified no RNA polymerase and TATA-box binding proteins in any of the ten complete Borg genomes, it is possible that *Methanoperedens* can tightly regulate Borg gene transcription, thus controlling the production and localization of Borg proteins. TRs may be involved in this process. Approximately half of the TR regions were intergenic or have the first unit within or partially within gene ends. These could be (long) noncoding RNAs (lncRNAs), which are known to modulate chromosome structure and function, transcription of neighboring and distant genes, affect RNA stability and translation and serve as architectural RNAs (arcRNAs) [29]. arcRNAs are of particular interest as they are involved in forming RNA-protein condensates, and some arcRNAs possess repeat sequences to accumulate multiple copies of specific proteins and/or multiple copies of RNAs [30].

Virtually all TRs in ORF were of lengths divisible by three, thus do not disrupt ORFs, and their A bias was not as strong as intergenic TRs. This indicates a very strong selective pressure to always introduce amino acid repeats and a selection for (or deselection against) specific codons. Bulky or highly reactive amino acids such as cysteine, which could severely interfere with redox homeostasis, were absent. Yet, disorder-promoting amino acids were enriched in aaTRs and concomitantly most aaTR regions were intrinsically disordered. Thus, we infer that aaTRs impact protein functioning. While protein homologues from *Methanoperedens* (or other microbial genomes) almost never showed aaTRs, Borg homologues (or functionally related Borg proteins) had aaTRs that were highly similar in aa character, but not identical in DNA sequence. For very closely related Borgs, this may arise because similar DNA regions in Borg genes were seeds for TR formation. However, the presence of aaTRs in different regions of proteins with the same function (e.g., MHCs), indicates a strong selective pressure for specific aaTRs that is constrained by protein function. Overall, we consider the combination of features to be strong evidence that the aaTRs are not random genomic aberrations and those in ORFs fulfill an important role in protein functioning.

To shed light on the predicted function that the proteins gain by having evolved aaTRs, we screened many of aaTRs for predicted functional sites. Most aaTRs are IDRs and these are commonly found in SLiMs, and vice versa [12]. In the case of the N-terminal aaTRs in the Borg ribonucleoproteins, the long loops that extend from the RNA-binding protein core could serve as docking sites for interaction partners. This is reminiscent of eukaryotic homologues which possess fused C-terminal IDR domains and are part of the spliceosome, a ribonucleoprotein machine that excises introns from eukaryotic pre-mRNAs [31]. The aaTR extension of the Borg ribonucleoproteins may undergo a disorder-to-order transition induced by binding to a yet to be identified protein partner, which could redefine its function in e.g., pre-mRNA or pre-tRNA processing [32]. The aaTRs associated with MHCs were also predicted to form long, largely unstructured extensions. Their consistent compositional dominance by proline and threonine clearly suggests a shared function and convergent evolution of these aaTRs. They could be intraprotein intersections that are glycosylated or phosphorylated, triggering a signaling cascade and conformational changes that lead to a modified electron flow, redox capacity and possibly nature of electron acceptor. The aaTRs could also be involved in navigating interprotein interfaces by being docking sites for other MHCs or extracellular oxidoreductases, such as a manganese oxidase that we found in Lilac Borg [3]. We thus propose that the Borg MHCs expand the redox and metabolic capacity of a large MHC network at the cell surface of the host [33] and aaTRs within them may enable a tunable connection to the electron conduction system that is integral to *Methanoperedens’* metabolism.

We found that aaTRs were statistically enriched in Borg proteins with predicted extracellular and membrane localization, another indication that their formation is not a random event. These proteins are typically implicated in cell-cell interactions, transport, protection and virulence. It is unclear whether Borgs have an existence outside of *Methanoperedens* cells, but the inferred high copy number of some Borgs compared to host cells [3] may point to this possibility. Borg-encoded S-layer and PEGA domain proteins (with and without aaTRs) could potentially encapsulate Borgs and mimic host proteins to evade host defense. Alternatively (or at a different time in their existence), these Borg proteins could be displayed on the *Methanoperedens* cell surface to alter host processes.

If Borgs can exist outside of host cells, they would need the ability to infect *Methanoperedens* cells to replicate (analogous to viruses). Proteins involved in infection and defense are expected to be fast evolving, and TRs may serve as a mechanism to enable this. A hotspot for TR evolution that is conserved across Borgs encodes TolA (and additional conserved hypothetical proteins). TolA is important for cell envelope integrity and is crucial for entry of filamentous bacteriophages that infect *Escherichia coli* or *Vibrio cholerae* [34,35]. TolA is exceptional, as unlike other Borg aaTR-proteins, homologous proteins from *Streptomyces* and *Bacillus* are also repeat proteins with similar aa signatures (EAKQ). The observation that the TolA-containing regions in Borgs are conserved, fast evolving and under strong selective pressure could be consistent with a role in Borg-host attachment. As proteins produced in the host they may facilitate cell-cell interaction (e.g., for lateral gene transfer).

A striking observation from prior work at the wetland site is the large diversity of different and coexisting Borgs and *Methanoperedens* species [3]. Based on correlation analyses, different Borgs associate with different *Methanoperedens* [3], implying co-evolution of Borgs and hosts. Lateral gene transfer has been reported as a driver for metabolic flexibility in members of the *Methanoperedenaceae* [36]. Gene acquisition (lateral gene transfer) is a prominent capacity of Borgs (this gave rise to their name) and may contribute to *Methanoperedens* speciation. TR regions also may be key for establishing new Borg-host associations, especially if the aaTRs and noncoding TRs enable functional cooperation between Borg and host protein inventories. Thus, while CRISPR repeats evolve rapidly to defend against phage infection, rapid evolution of Borg tandem repeats may be required to maintain coexistence with their hosts during coevolution.

## Conclusion

We conclude that the nucleotide regions flanking repeats, and the uniqueness of the tandem repeats in each locus likely indicate that TRs arise from local sequences rather than being introduced from external templates. Other constraints on the mechanism behind TR formation are their generally perfect sequence repetition, and A-enriched composition. Many tandem repeats lead to aaTRs in proteins, and these are usually intrinsically disordered regions that are also predicted post-translational modification sites, and protein or nucleic acid binding sites. We propose that aaTR-proteins expand and modify the cellular and metabolic capacity of Borg-bearing *Methanoperedens*, yet expression of their large gene inventories is likely under tight control. Introduction of TRs in both genes and intergenic regions may be central to regulating Borg gene expression, translation, protein localization and function. TR regions change rapidly in number and distribution to generate within population heterogeneity and between population diversity. This feature is likely central for Borg infection, association and co-speciation of Borgs and their *Methanoperedens* hosts.

## Methods

### Identification of Borg genomes and manual genome curation

Metagenomic datasets on ggKbase (ggkbase.berkeley.edu) were searched for contigs with a dominant taxonomic profile matching *Methanoperedens* (Archaea; Euryarchaeaota; Methanomicrobia; Methanosarcinales; Candidatus Methanoperedens; Candidatus Methanoperedens nitroreducens). Manual genome binning was performed based on contig taxonomy, coverage, GC content (25-35% GC) and presence of nucleotide tandem repeats.

Manual curation of Borg genomes was performed in Geneious Prime 2021.2.2 (https://www.geneious.com). It involved piecing together, and verifying by paired reads placement, fragments of approximately the same GC content, sequencing reads coverage, phylogenetic profile and relatedness to known Borgs into a single chromosome. Subsequently, careful visualization of the patterns of read discrepancies was used to locate local assembly errors, most of which were fixed by either relocating paired reads or introducing previously unplaced paired reads.

### Genome visualization, alignments and GC skew analysis

Genomes were visualized in Geneious Prime 2021.2.2 and aligned with the MCM algorithm, or the progressiveMauve algorithm when aligning multiple contigs. Genetic neighborhood comparisons were performed using clinker [37] (v0.0.21).

GC skew and cumulative GC skew were calculated as described previously [38].

### Tandem repeat identification

Nucleotide tandem repeats were initially predicted in Geneious Prime 2021.2.2 (Repeat Finder, minimum repeat length 50, maximum mismatches 0) and then marked down in the completed genomes using a Python script (https://github.com/rohansachdeva/tandem_repeats) based on MUMmer (v3.23) [39]. Nucleotide tandem repeats were searched using a stringent threshold of ≥ 50 nt region and ≥ 3 TR units and no mismatch (--min_region_len 50 --min_repeat_count 3 -p 10). Amino acid tandem repeats were searched using a stringent threshold of ≥ 16 aa and ≥ 3 TR (-l 3 --min_repeat_count 3 --min_uniq_nt 1 --min_region_len 16 -p 30).

### Secondary structure prediction

The secondary structure of tandem repeats was predicted with RNAfold WebServer [40].

### Protein family clustering

A dataset of 11,995 Borg proteins was constructed using all proteins from the ten curated Borg genomes and 37 additional aaTR-proteins from curated contigs of Pink, Blue, Steel, Olive, Grey and Apricot Borg. All proteins were clustered into protein subfamilies using MMseqs [41] and HMMs were constructed from these subfamilies using HHblits [42] as previously described [43]. They were then profiled against the PFAM database by HMM-HMM comparison using HHsearch [44] and protein subfamilies enriched in plasmid proteins were determined as described previously [45].

### Functional prediction of aaTR-proteins

aaTR-proteins were profiled using InterProScan [46] (v5.51-85.0) to get functional and structural annotations of individual proteins. Protein subfamilies were functionally annotated using HMMER (hmmer.org) (v3.3, hmmsearch) and the PFAM (--cut_nc) HMM database [47]. Homology search was performed with blastp [48]. Intrinsic disorder was predicted with MobiDBLite [49] and IUPred2A [50], transmembrane helices and cellular localization were predicted with TMHMM [51] (v2.0) and psort [52] (v2.0, archaeal mode). SLiMs were predicted with the eukaryotic linear motif (ELM) resource for functional sites in proteins [14] (http://elm.eu.org/). The single aaTR unit was queried (cell compartment: not specified) and the functional site covering most of the aaTR was usually selected (Table S3). Functions of aaTR regions in proteins were also predicted with flDPnn [53].

### Hierarchical clustering of aaTRs and construction of a phylogenetic tree

Hierarchical clustering of aaTRs was performed using triple tandem repeat units for the alignments, since three was the minimum repeat unit length to make the threshold and the regions were dynamic. The alignments were performed with MAFFT (--treeout --reorder --localpair) [54] (v7.453). The aaTR clusters were visualized and decorated in iTOL [55]. An aaTR cluster was formed when a branch contained 4≥ related sequences. The names of the clusters were given based on the most abundant amino acids found in the repeat units (amino acids represented by 10%≥ became name-giving).

The Borg DNA polymerase sequences were aligned with a reference dataset from [11], aligned with MAFFT (--reorder --auto) [54] (v7.453), trimmed with trimal (-gt 0.2) [56] (v1.4.rev15) and a maximum-likelihood tree was calculated in IQ-TREE (-m TEST -st AA -bb 1000 -nt AUTO -ntmax 20 -pre) [57]. The phylogenetic tree of the DNA polymerases was visualized and decorated in iTOL [55].

### Structural modeling

Structural modeling of aaTR-bearing Sm proteins was initially performed using SWISS-MODEL [58] and the best hit in the Swiss-Prot database as template. Further structural modeling of aaTR-bearing Sm proteins was performed with AlphaFold2 using ColabFold [59] and selecting the experimental option homooligomer 6. Structural modeling of aaTR-MHC proteins was performed using AlphaFold2 [60] via a LocalColabFold (--use_ptm --use_turbo --num_relax Top5 --max_recycle 3) [61,62]. Modeled protein structures were visualized and superimposed onto PDB structures using PyMOL [63] (v2.3.4).

## Acknowledgements

This publication is based on research in part funded by the Bill & Melinda Gates Foundation. The findings and conclusions contained within are those of the authors and do not necessarily reflect positions or policies of the Bill & Melinda Gates Foundation. Funding for this research was also provided by a DFG fellowship for MCS (Project Number: 447383558), and by the Innovative Genomics Institute at UC Berkeley. We thank Shufei Lei, Jordan Hoff and Adair Borges for bioinformatics assistance and Luis Valentin Alvarado and Susan Marquesee for helpful discussions.

## Author contributions

MCS and JFB designed the study. JFB performed the binning and carried out the manual genome curation. MCS performed genome alignments, TR analyses, secondary structure predictions, protein family clustering, hierarchical clustering and phylogenetic tree construction, protein analyses, and structural modeling with the help of RS, LW, JWR and JFB. RS wrote the code to detect TRs. MCS and JFB did GC skew analyses and interpreted the results. MCS and JFB wrote the manuscript with input from all authors.

## Data availability

Newly released Borg genomes used in this manuscript are available via: https://ggkbase.berkeley.edu/tandem_repeats_in_borgs. The publicly available dataset used for other *Methanoperedens* analyses can be accessed via: https://ggkbase.berkeley.edu/BMp/organisms. Protein sequences, structural models and the phylogenetic tree of the DNA polymerases from this study are available through Zenodo (10.5281/zenodo.6533809). The python script that was used to detect tandem repeats on the nucleotide or protein level is available on GitHub (https://github.com/rohansachdeva/tandem_repeats). Statistics on TR regions in the ten completed Borg genomes can be found in Supplementary Table S8 and Table S9.

## Supplementary Figures

**Figure S1.**
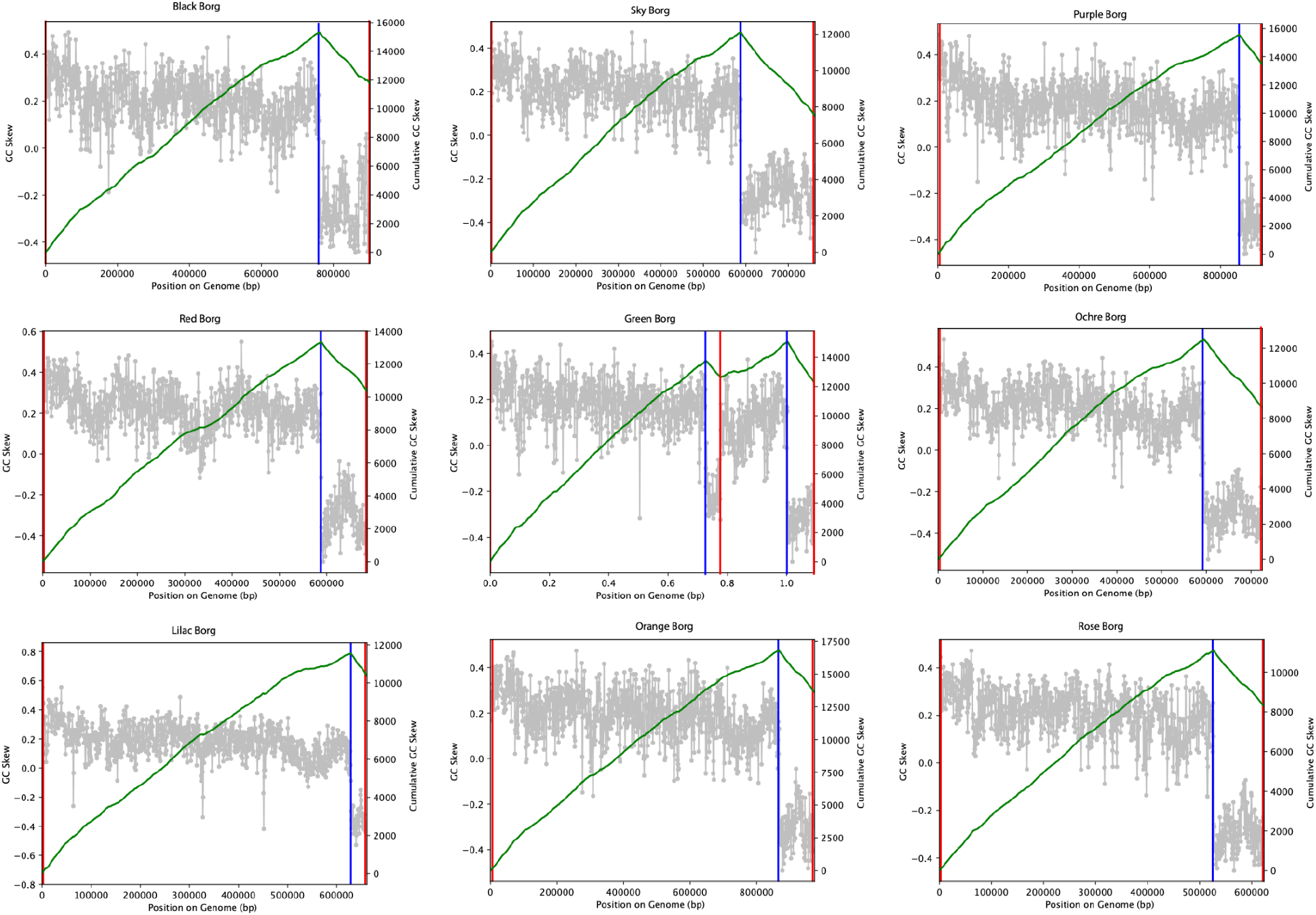
Borg genome replication is initiated at the termini. Shown is the GC skew (gray) and cumulative GC skew (green lines). Borg DNA is replicated from the terminal inverted repeats (origin, red lines) until the terminus (terminus, blue lines).

**Figure S2.**
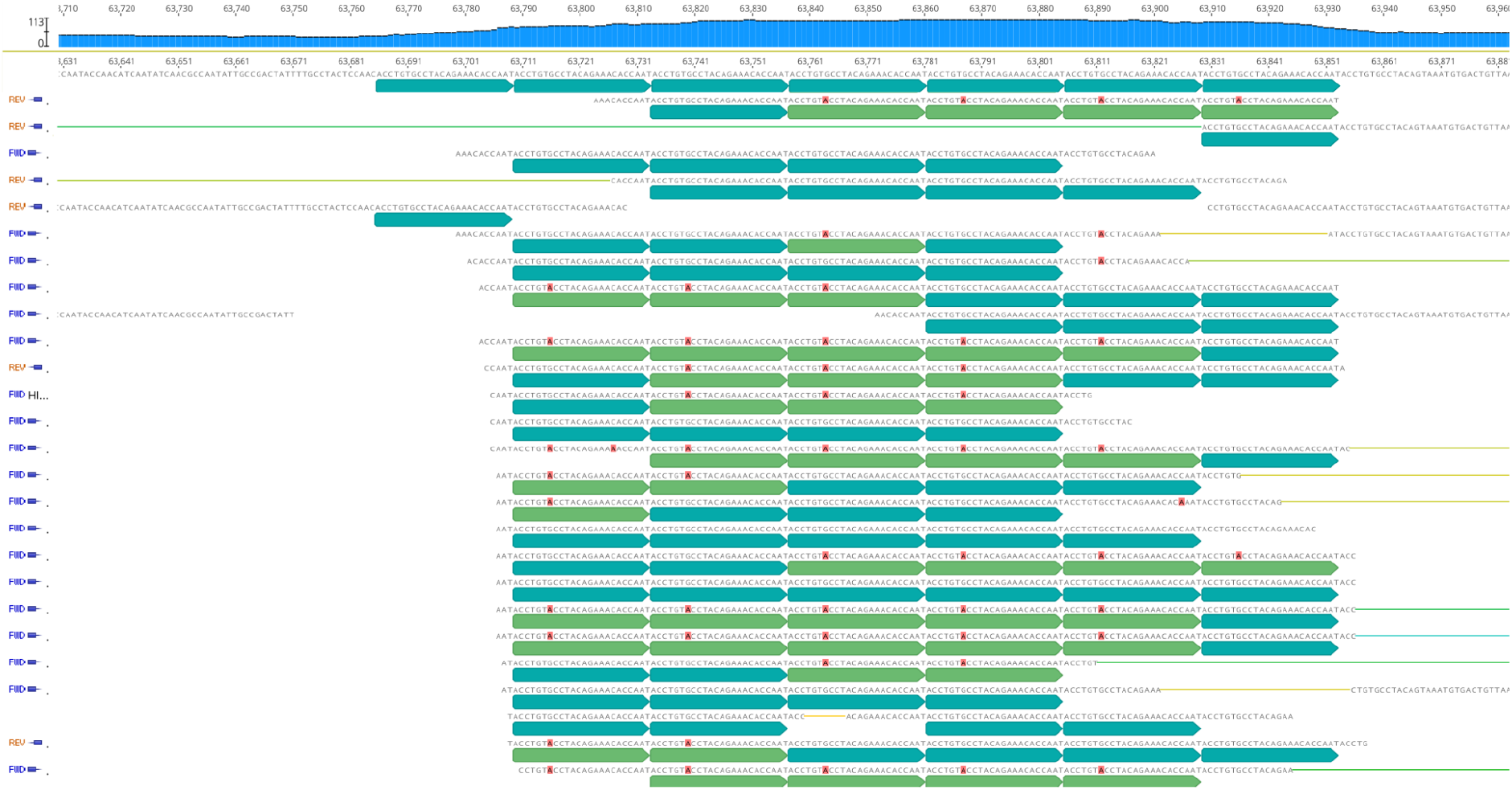
Reads mapped to a *de novo* assembly showing different combinations of repeat units in a Borg subpopulation. There are two types of repeat units (magenta or green segments). They are identical, with the exception of a SNP (highlighted in red).

**Figure S3.**
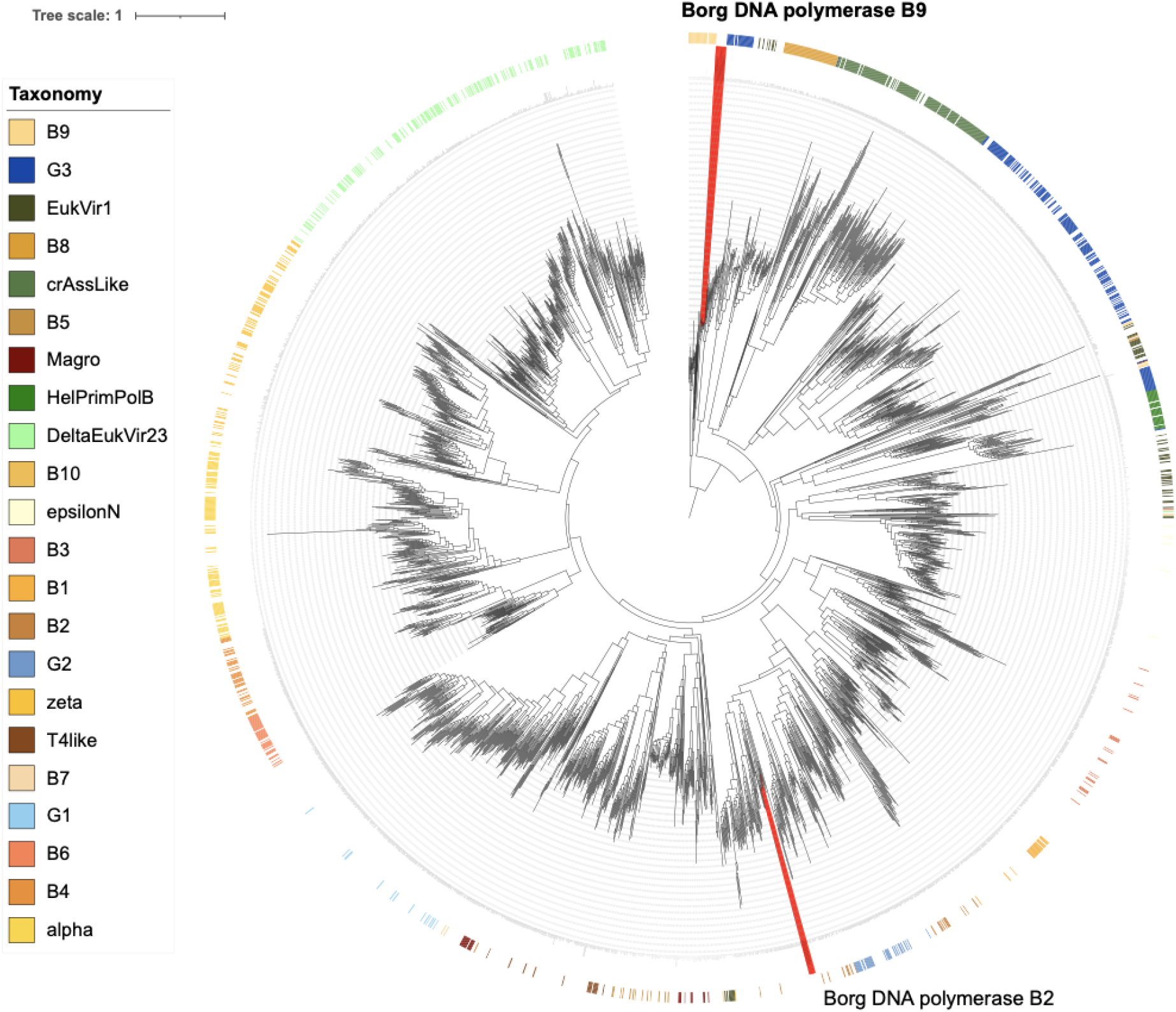
Phylogenetic placement of Borg DNA polymerases. Amino acid sequences of marker DNA polymerases cluster together in the B9 clade. Additional DNA polymerases present in some Borgs cluster together in the B2 clade. Reference sequences originate from Kazlauskas *et al*. (2020). The tree was rooted between the G3 and B9 clade.

**Figure S4.**
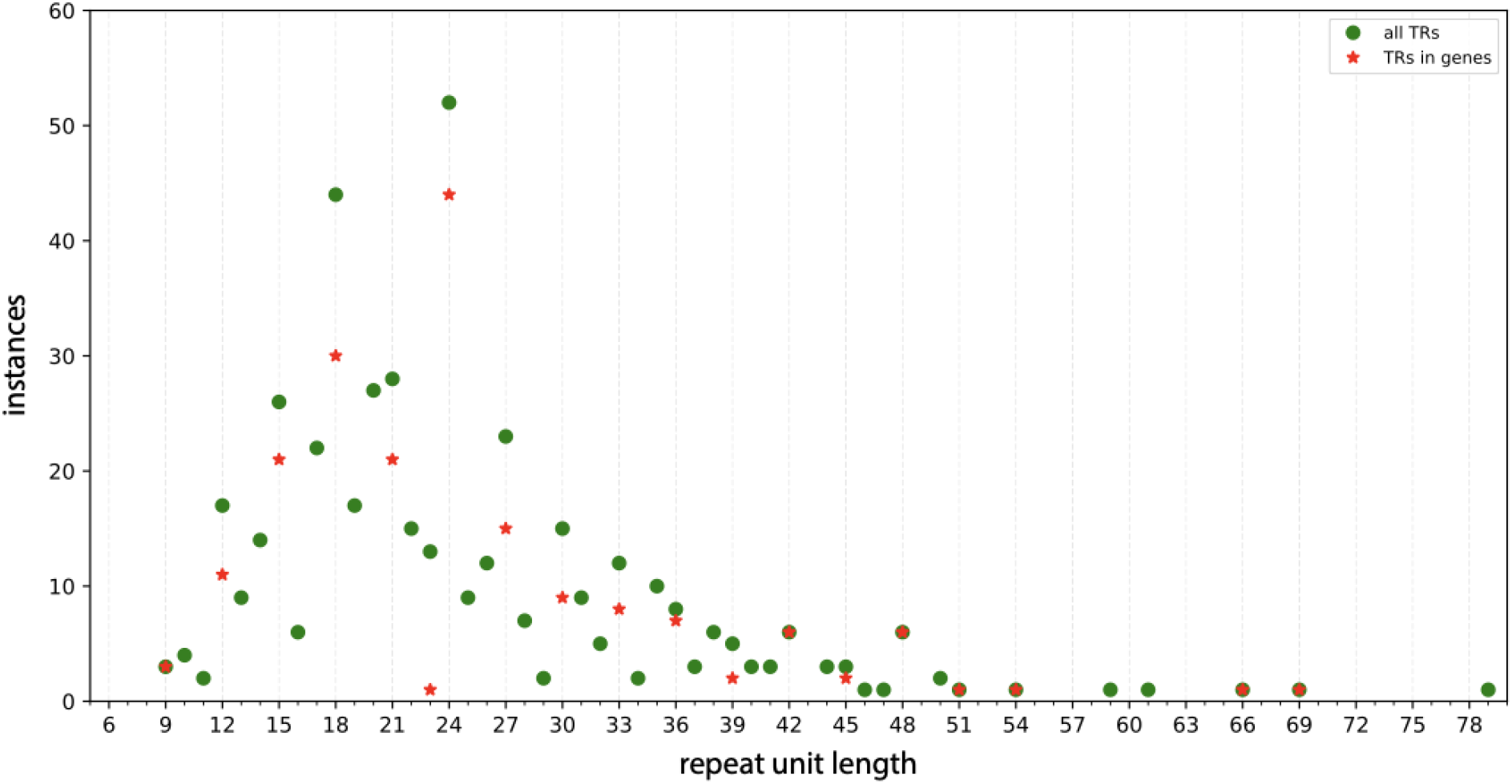
Tandem repeat unit lengths of all 460 TR regions across 10 Borgs.

**Figure S5.**
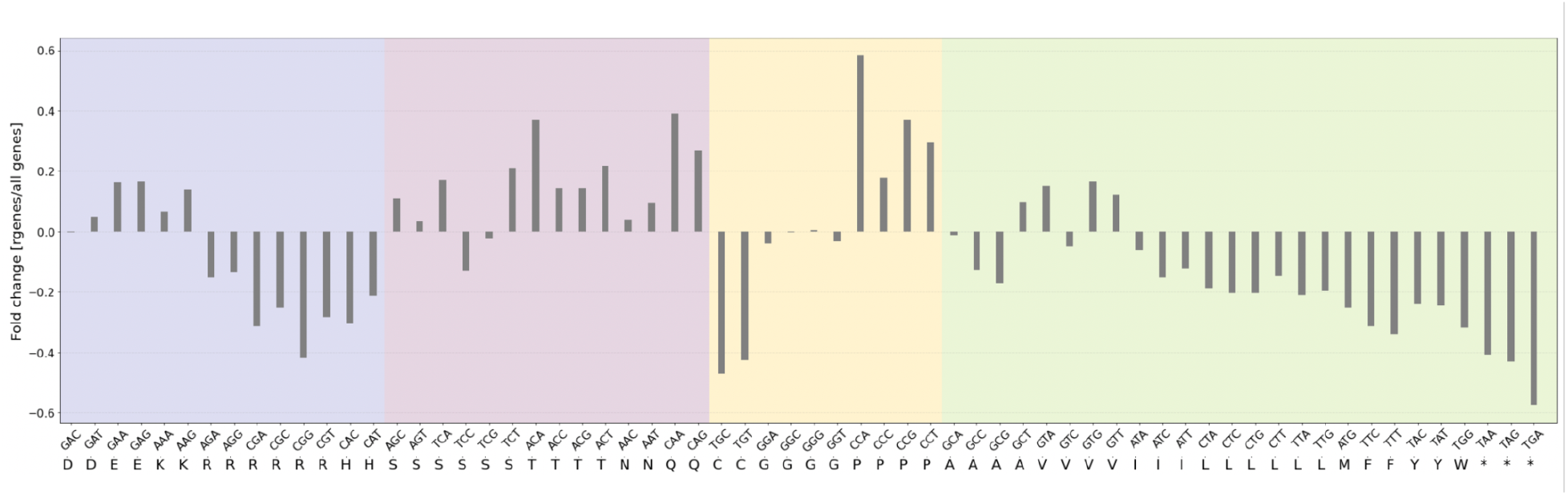
Relative codon frequency in TR-bearing ORFs versus all ORFs. The codon frequency (frequency/thousand) of ORFs with TRs (rgenes) was divided by the codon frequency of all ORFs from all ten complete Borg genomes, and plotted for each codon and its corresponding amino acid. Codons more frequent in ORFs with TRs compared to all Borg ORFs show a positive fold change, codons less frequent show a negative fold change.

**Figure S6.**
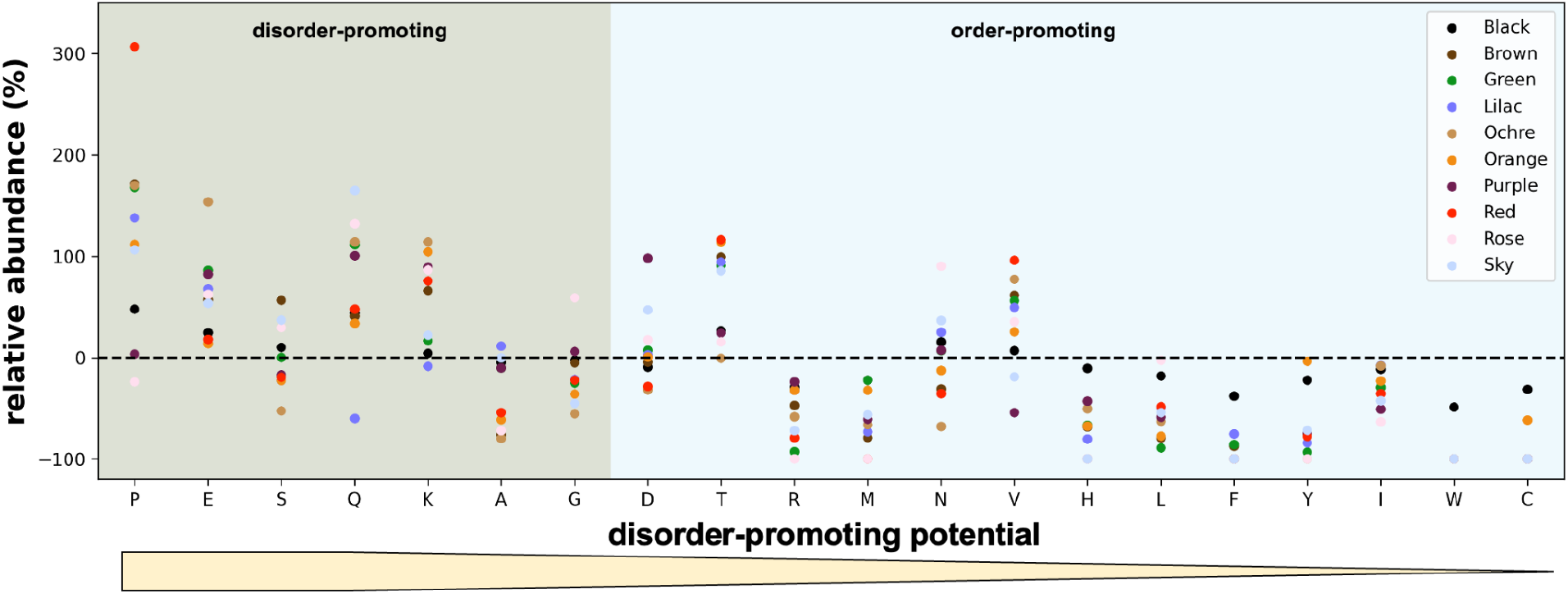
aaTRs are enriched in disorder-promoting amino acids. The aa abundance reflects the frequency of amino acids in aaTRs divided by the frequency in all proteins.

**Figure S7.**
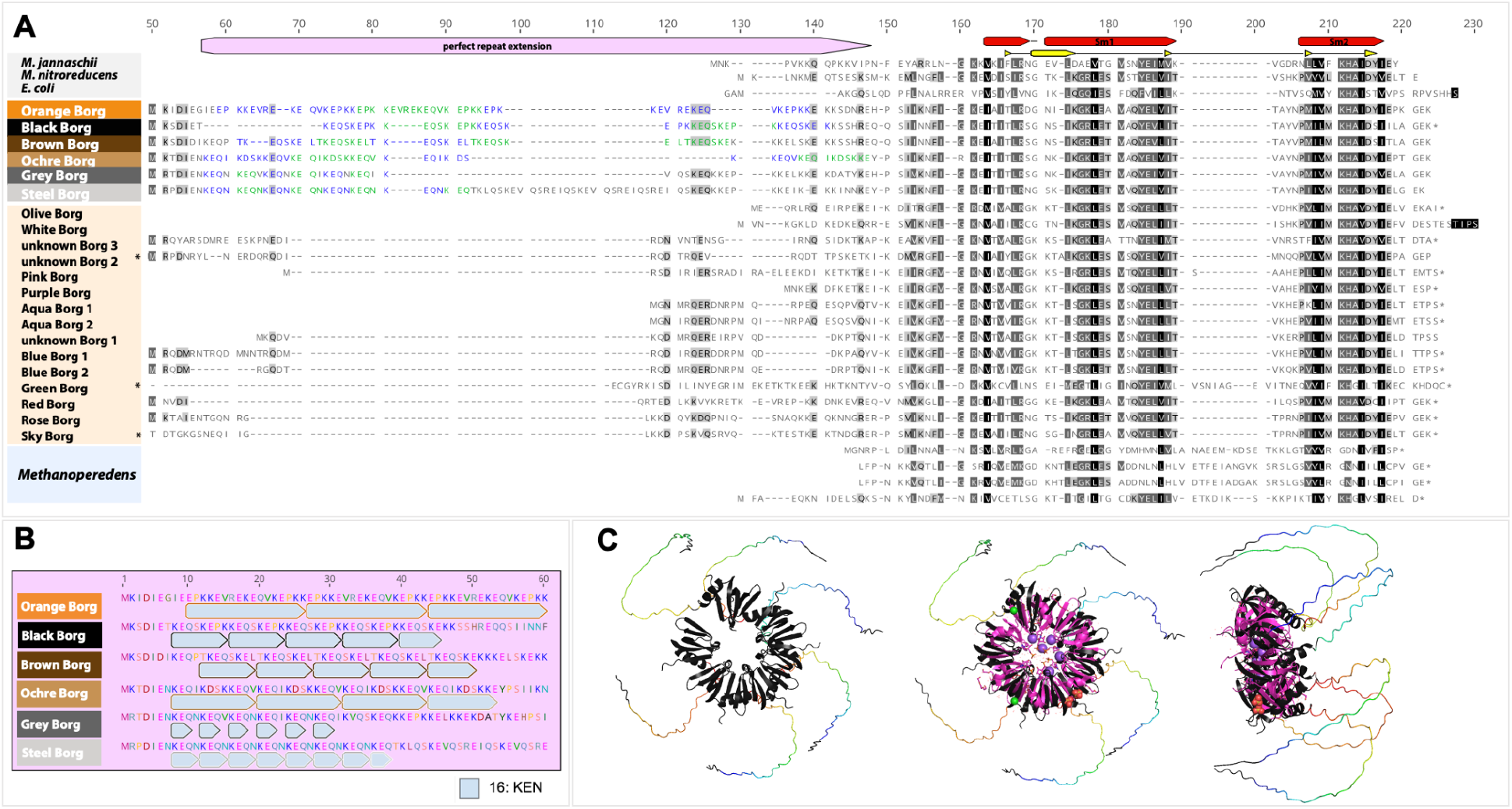
Sequence alignment, aaTR composition and predicted structures of Borg Sm ribonucleoproteins. **A**. Multiple sequence alignment of Borg Sm ribonucleoproteins with and without aaTRs, Sm from *Methanoperedens* bins co-occurring with Borgs and reference sequences from *M. jannaschii* (PDB: 4×9D), *M. nitroreducens* (WP_096203417) and *E. coli* (PDB: 1HK9). **B**. aaTR units of Sm ribonucleoproteins. **C**. Predicted structures of aaTR-Sm ribonucleoproteins.

## Notes

### Competing Interest Statement

J.F.B. is a founder of Metagenomi.

## References

1. Lai S, Jia L, Subramanian B, Pan S, Zhang J, Dong Y, et al. mMGE: a database for human metagenomic extrachromosomal mobile genetic elements. Nucleic Acids Res. 2021;49: D783–D791.

2. Yu MK, Fogarty EC, Murat Eren A. The genetic and ecological landscape of plasmids in the human gut. bioRxiv. 2022. p. 2020.11.01.361691. doi:10.1101/2020.11.01.361691

3. Al-Shayeb B, Schoelmerich MC, West-Roberts J, Valentin-Alvarado LE, Sachdeva R, Mullen S, et al. Borgs are giant extrachromosomal elements with the potential to augment methane oxidation. bioRxiv. 2021. p. 2021.07.10.451761. doi:10.1101/2021.07.10.451761.Please see the updated version provided with the manuscript files. Unfortunately, the journal prohibits an upload of the current version. We anticipate that the paper will be “in press” very shortly.

4. Schoelmerich MC, Oubouter HT, Sachdeva R, Penev P, Amano Y, West-Roberts J, et al. A widespread group of large plasmids in methanotrophic Methanoperedens archaea. bioRxiv. 2022. p. 2022.02.01.478723. doi:10.1101/2022.02.01.478723

5. Haroon MF, Hu S, Shi Y, Imelfort M, Keller J, Hugenholtz P, et al. Anaerobic oxidation of methane coupled to nitrate reduction in a novel archaeal lineage. Nature. 2013;500: 567–570.

6. Koonin EV, Yutin N. Evolution of the Large Nucleocytoplasmic DNA Viruses of Eukaryotes and Convergent Origins of Viral Gigantism. Adv Virus Res. 2019;103: 167–202.

7. Wang H, Peng N, Shah SA, Huang L, She Q. Archaeal extrachromosomal genetic elements. Microbiol Mol Biol Rev. 2015;79: 117–152.

8. Gunge N, Fukuda K, Takahashi S, Meinhardt F. Migration of the yeast linear DNA plasmid from the cytoplasm into the nucleus in Saccharomyces cerevisiae. Curr Genet. 1995;28: 280–288.

9. Chater KF, Kinashi H. Streptomyces Linear Plasmids: Their Discovery, Functions, Interactions with Other Replicons, and Evolutionary Significance. In: Meinhardt F, Klassen R, editors. Microbial Linear Plasmids. Berlin, Heidelberg: Springer Berlin Heidelberg; 2007. pp. 1–31.

10. Wagenknecht M, Dib JR, Thürmer A, Daniel R, Farías ME, Meinhardt F. Structural peculiarities of linear megaplasmid, pLMA1, from Micrococcus luteus interfere with pyrosequencing reads assembly. Biotechnol Lett. 2010;32: 1853–1862.

11. Kazlauskas D, Krupovic M, Guglielmini J, Forterre P, Venclovas Č. Diversity and evolution of B-family DNA polymerases. Nucleic Acids Res. 2020;48: 10142–10156.

12. van der Lee R, Buljan M, Lang B, Weatheritt RJ, Daughdrill GW, Dunker AK, et al. Classification of intrinsically disordered regions and proteins. Chem Rev. 2014;114: 6589–6631.

13. Van Roey K, Uyar B, Weatheritt RJ, Dinkel H, Seiler M, Budd A, et al. Short linear motifs: ubiquitous and functionally diverse protein interaction modules directing cell regulation. Chem Rev. 2014;114: 6733–6778.

14. Kumar M, Gouw M, Michael S, Sámano-Sánchez H, Pancsa R, Glavina J, et al. ELM-the eukaryotic linear motif resource in 2020. Nucleic Acids Res. 2020;48: D296–D306.

15. Maupin-Furlow J. Proteasomes and protein conjugation across domains of life. Nat Rev Microbiol. 2011;10: 100–111.

16. Garnham CP, Roll-Mecak A. The chemical complexity of cellular microtubules: tubulin post-translational modification enzymes and their roles in tuning microtubule functions. Cytoskeleton. 2012;69: 442–463.

17. Vogel J, Luisi BF. Hfq and its constellation of RNA. Nat Rev Microbiol. 2011;9: 578–589.

18. Nikulin A, Mikhailina A, Lekontseva N, Balobanov V, Nikonova E, Tishchenko S. Characterization of RNA-binding properties of the archaeal Hfq-like protein from Methanococcus jannaschii. J Biomol Struct Dyn. 2017;35: 1615–1628.

19. Kletzin A, Heimerl T, Flechsler J, van Niftrik L, Rachel R, Klingl A. Cytochromes c in Archaea: distribution, maturation, cell architecture, and the special case of Ignicoccus hospitalis. Front Microbiol. 2015;6: 439.

20. Ryan CP. Tandem repeat disorders. Evol Med Public Health. 2019;2019: 17.

21. Usdin K. The biological effects of simple tandem repeats: lessons from the repeat expansion diseases. Genome Res. 2008;18: 1011–1019.

22. Fondon JW 3rd, Garner HR. Molecular origins of rapid and continuous morphological evolution. Proc Natl Acad Sci U S A. 2004;101: 18058–18063.

23. Viguera E, Canceill D, Ehrlich SD. Replication slippage involves DNA polymerase pausing and dissociation. EMBO J. 2001;20: 2587–2595.

24. Zhou K, Aertsen A, Michiels CW. The role of variable DNA tandem repeats in bacterial adaptation. FEMS Microbiol Rev. 2014;38: 119–141.

25. Kim JC, Mirkin SM. The balancing act of DNA repeat expansions. Curr Opin Genet Dev. 2013;23: 280–288.

26. Castillo-Lizardo M, Henneke G, Viguera E. Replication slippage of the thermophilic DNA polymerases B and D from the Euryarchaeota Pyrococcus abyssi. Front Microbiol. 2014;5: 403.

27. Tyson GW, Banfield JF. Rapidly evolving CRISPRs implicated in acquired resistance of microorganisms to viruses. Environ Microbiol. 2008;10: 200–207.

28. McGinn J, Marraffini LA. Molecular mechanisms of CRISPR-Cas spacer acquisition. Nat Rev Microbiol. 2019;17: 7–12.

29. Statello L, Guo C-J, Chen L-L, Huarte M. Gene regulation by long non-coding RNAs and its biological functions. Nat Rev Mol Cell Biol. 2021;22: 96–118.

30. Ninomiya K, Hirose T. Short Tandem Repeat-Enriched Architectural RNAs in Nuclear Bodies: Functions and Associated Diseases. Noncoding RNA. 2020;6. doi:10.3390/ncrna6010006

31. Coelho Ribeiro M de L, Espinosa J, Islam S, Martinez O, Thanki JJ, Mazariegos S, et al. Malleable ribonucleoprotein machine: protein intrinsic disorder in the Saccharomyces cerevisiae spliceosome. PeerJ. 2013;1: e2.

32. Törö I, Thore S, Mayer C, Basquin J, Séraphin B, Suck D. RNA binding in an Sm core domain: X-ray structure and functional analysis of an archaeal Sm protein complex. EMBO J. 2001;20: 2293–2303.

33. Breuer M, Rosso KM, Blumberger J. Electron flow in multiheme bacterial cytochromes is a balancing act between heme electronic interaction and redox potentials. Proc Natl Acad Sci U S A. 2014;111: 611–616.

34. Yakhnina AA, Bernhardt TG. The Tol-Pal system is required for peptidoglycan-cleaving enzymes to complete bacterial cell division. Proc Natl Acad Sci U S A. 2020;117: 6777–6783.

35. Heilpern AJ, Waldor MK. CTXphi infection of Vibrio cholerae requires the tolQRA gene products. J Bacteriol. 2000;182: 1739–1747.

36. Leu AO, McIlroy SJ, Ye J, Parks DH, Orphan VJ, Tyson GW. Lateral Gene Transfer Drives Metabolic Flexibility in the Anaerobic Methane-Oxidizing Archaeal Family Methanoperedenaceae. MBio. 2020;11. doi:10.1128/mBio.01325-20

37. Gilchrist CLM, Chooi Y-H. Clinker & clustermap.js: Automatic generation of gene cluster comparison figures. Bioinformatics. 2021. doi:10.1093/bioinformatics/btab007

38. Brown CT, Olm MR, Thomas BC, Banfield JF. Measurement of bacterial replication rates in microbial communities. Nat Biotechnol. 2016;34: 1256–1263.

39. Kurtz S, Phillippy A, Delcher AL, Smoot M, Shumway M, Antonescu C, et al. Versatile and open software for comparing large genomes. Genome Biol. 2004;5: R12.

40. Gruber AR, Lorenz R, Bernhart SH, Neuböck R, Hofacker IL. The Vienna RNA websuite. Nucleic Acids Res. 2008;36: W70–4.

41. Hauser M, Steinegger M, Söding J. MMseqs software suite for fast and deep clustering and searching of large protein sequence sets. Bioinformatics. 2016;32: 1323–1330.

42. Remmert M, Biegert A, Hauser A, Söding J. HHblits: lightning-fast iterative protein sequence searching by HMM-HMM alignment. Nat Methods. 2011;9: 173–175.

43. Méheust R, Burstein D, Castelle CJ, Banfield JF. The distinction of CPR bacteria from other bacteria based on protein family content. Nat Commun. 2019;10: 4173.

44. Söding J. Protein homology detection by HMM-HMM comparison. Bioinformatics. 2005;21: 951–960.

45. Jaffe AL, Thomas AD, He C, Keren R, Valentin-Alvarado LE, Munk P, et al. Patterns of Gene Content and Co-occurrence Constrain the Evolutionary Path toward Animal Association in Candidate Phyla Radiation Bacteria. MBio. 2021;12: e0052121.

46. Jones P, Binns D, Chang H-Y, Fraser M, Li W, McAnulla C, et al. InterProScan 5: genome-scale protein function classification. Bioinformatics. 2014;30: 1236–1240.

47. Finn RD, Bateman A, Clements J, Coggill P, Eberhardt RY, Eddy SR, et al. Pfam: the protein families database. Nucleic Acids Res. 2014;42: D222–30.

48. Altschul SF, Gish W, Miller W, Myers EW, Lipman DJ. Basic local alignment search tool. J Mol Biol. 1990;215: 403–410.

49. Necci M, Piovesan D, Dosztányi Z, Tosatto SCE. MobiDB-lite: fast and highly specific consensus prediction of intrinsic disorder in proteins. Bioinformatics. 2017;33: 1402–1404.

50. Erdős G, Dosztányi Z. Analyzing Protein Disorder with IUPred2A. Curr Protoc Bioinformatics. 2020;70: e99.

51. Krogh A, Larsson B, von Heijne G, Sonnhammer EL. Predicting transmembrane protein topology with a hidden Markov model: application to complete genomes. J Mol Biol. 2001;305: 567–580.

52. Yu NY, Wagner JR, Laird MR, Melli G, Rey S, Lo R, et al. PSORTb 3.0: improved protein subcellular localization prediction with refined localization subcategories and predictive capabilities for all prokaryotes. Bioinformatics. 2010;26: 1608–1615.

53. Hu G, Katuwawala A, Wang K, Wu Z, Ghadermarzi S, Gao J, et al. flDPnn: Accurate intrinsic disorder prediction with putative propensities of disorder functions. Nat Commun. 2021;12: 4438.

54. Katoh K, Misawa K, Kuma K-I, Miyata T. MAFFT: a novel method for rapid multiple sequence alignment based on fast Fourier transform. Nucleic Acids Res. 2002;30: 3059–3066.

55. Letunic I, Bork P. Interactive tree of life (iTOL) v3: an online tool for the display and annotation of phylogenetic and other trees. Nucleic Acids Res. 2016;44: W242–5.

56. Capella-Gutiérrez S, Silla-Martínez JM, Gabaldón T. trimAl: a tool for automated alignment trimming in large-scale phylogenetic analyses. Bioinformatics. 2009;25: 1972–1973.

57. Nguyen L-T, Schmidt HA, von Haeseler A, Minh BQ. IQ-TREE: a fast and effective stochastic algorithm for estimating maximum-likelihood phylogenies. Mol Biol Evol. 2015;32: 268–274.

58. Waterhouse A, Bertoni M, Bienert S, Studer G, Tauriello G, Gumienny R, et al. SWISS-MODEL: homology modelling of protein structures and complexes. Nucleic Acids Res. 2018;46: W296–W303.

59. Mirdita M, Ovchinnikov S, Steinegger M. ColabFold - Making protein folding accessible to all. bioRxiv. 2021. p. 2021.08.15.456425. doi:10.1101/2021.08.15.456425

60. Jumper J, Evans R, Pritzel A, Green T, Figurnov M, Ronneberger O, et al. Highly accurate protein structure prediction with AlphaFold. Nature. 2021;596: 583–589.

61. Mirdita M, Schütze K, Moriwaki Y, Heo L, Ovchinnikov S, Steinegger M. ColabFold - Making protein folding accessible to all. Research Square. 2021. doi:10.21203/rs.3.rs-1032816/v1

62. Moriwaki Y. localcolabfold: ColabFold on your local PC. Github; Available: https://github.com/YoshitakaMo/localcolabfold

63. DELANO, W. L. The PyMOL Molecular Graphics System. http://www.pymol.org. 2002 x[cited 27 Jan 2022]. Available: https://ci.nii.ac.jp/naid/10020095229/

